# Genome-wide analysis of Respiratory burst oxidase homolog (*Rboh*) genes in *Aquilaria* species and its association with agarwood formation

**DOI:** 10.1101/2023.05.08.539809

**Authors:** Khaleda Begum, Ankur Das, Raja Ahmed, Suraiya Akhtar, Ram Kulkarni, Sofia Banu

**Affiliations:** Department of Bioengineering and Technology, Gauhati University, Guwahati, Assam, India, 781014; Symbiosis School of Biological Sciences, Symbiosis International (Deemed University), Lavale, Pune, India

**Keywords:** *Aquilaria*, *Rboh*, ROS, H_2_O_2_, qRT-PCR

## Abstract

Respiratory burst oxidase homolog (*Rboh*), generate reactive oxygen species (ROS) to maintain normal growth and pathogen induced defence responses in plants. In *Aquilaria* plants, wounding and fungal invasion results in the biosynthesis of secondary metabolites as a defence response which with due course develop into agarwood. During pathogen invasion, *Aquilaria* tree accumulate ROS species through the action of Rboh enzymes. Although in agarwood formation role of Rboh gene family has been implicated, an comprehensives study on *Rboh* gene family and information of its role during agarwood formation in missing. In this study, seven *Rboh* genes were identified from the genomes of two *Aquilaria* species viz., *Aquilaria agallocha* and *Aquilaria sinensis* and phylogenetically classified into five groups.

Stress response, hormone regulation, and development related regulatory elements were identified in the promoter regions. The protein sequences comprised of four conserved domains, an EF-hand domain, and a transmembrane region which they probably utilise for MAPK signaling, plant-pathogen interaction and plant hormone signal transduction pathways. Expression analyses revealed that among the seven members, *AaRbohA* and *AaRhobC* were involved in generation of ROS species, and also probably play role in agarwood formation. These findings provide valuable information regarding the Rboh members of *A. agallocha* which can be further used for functional analyses for in-depth understanding of ROS mediated signalling and regulation of agarwood formation.

**Graphical abstract:** 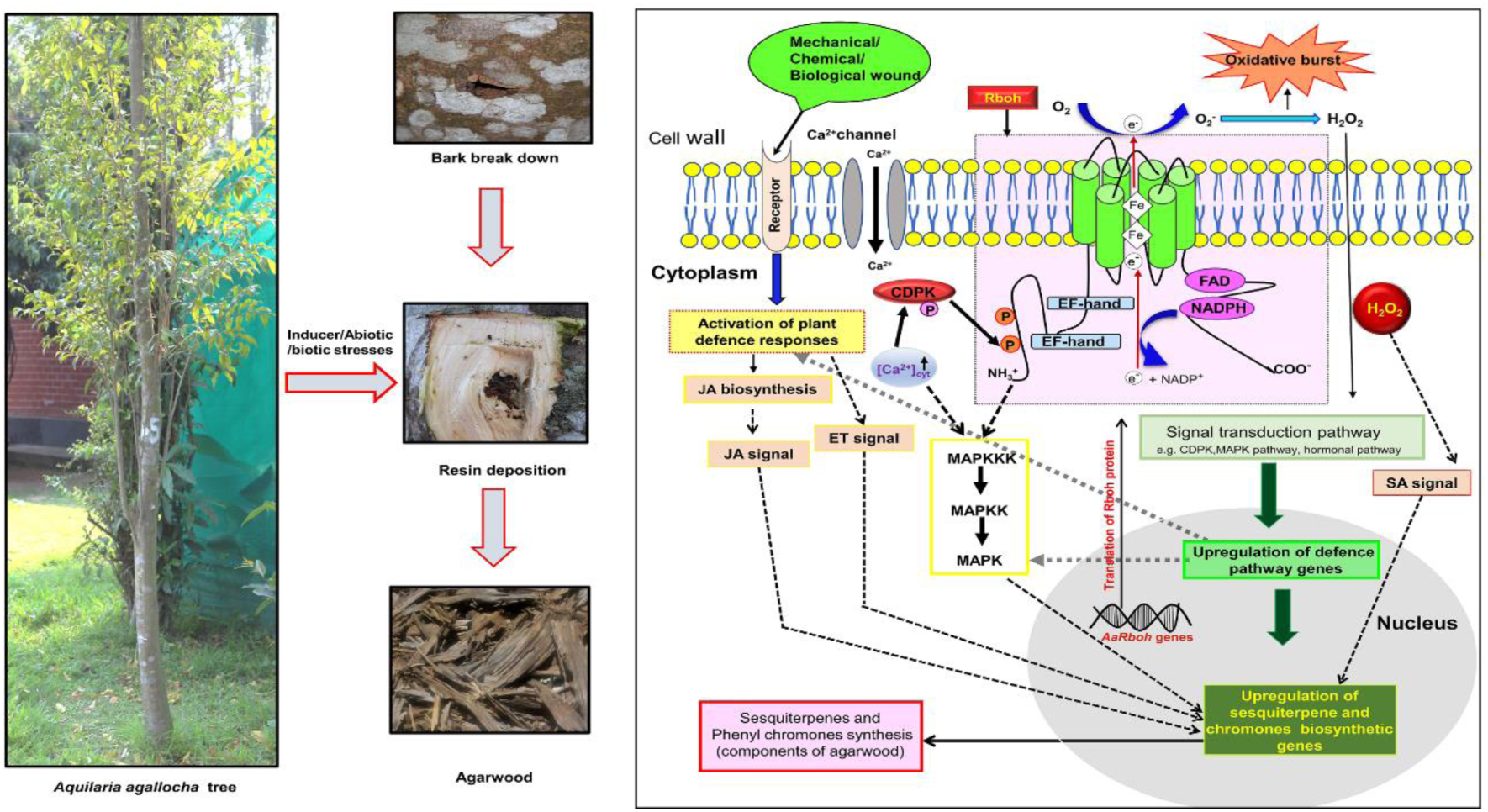

## 1. Introduction

The evolution of plants as sessile organisms creates challenges for them to cope continually with exposed environmental biotic and abiotic stresses, which tremendously affect their growths and yields (Wang et al., 2020; Mahilingam et al., 2021). To resist such stressors, including pathogenic infection, drought, and extreme temperature and salt concentrations, plant cells promote the accumulation of reactive oxygen species (ROS) and regulate nearly all biological processes associated with developmental stages, stresses and immunity responses (Castro et al., 2021). Although ROS was once considered as toxic by-products of aerobic metabolism (Inupakutika et al., 2016), recent studies affirm that it acts as a key signalling molecules to activate signal transduction cascades related to stress responses (Hu et al., 2018; Hawamda et al., 2020). However, beyond a specific threshold level, accumulation of ROS species can causes abnormal and irreparable metabolic changes and cell damages (Liu et al., 2016; Cheng et al., 2020; Wang et al., 2020). In plant, Hydrogen peroxide (H_2_O_2_) is the primary ROS species of peroxisomes produced during the C2 cycle (photorespiration) (Liu et al., 2016). Additionally, the photosynthesizing chloroplast and respiring mitochondria also produce superoxide and hydrogen peroxide as a by-product of photosynthesis and respiration (Navathe et al., 2019). The apoplastic molecular oxygen (O_2_) is first converted into superoxide anion (O^−^_2_) by Respiratory burst oxidase homolog protein (*Rboh*), which is converted to hydrogen peroxide (H_2_O_2_) through superoxide dismutation reaction by Superoxide dismutase (SOD, EC 1.15.1.1) (Navathe et al., 2019). The different *Rboh* genes or nicotinamide adenine dinucleotide phosphate (NADPH oxidase) in the plasma membrane transfer electrons from cytosolic NADPH/NADH to apoplastic oxygen, producing the reactive oxygen species in the cells (Yu et al., 2020). Structurally, plant Rboh proteins are intrinsic proteins having conserved six transmembrane α-helices bearing two basic helix– loop–helix calcium-binding structural domains (EF-hands) that are directly regulated by Ca^2+^ ions (Yu et al., 2020). Therefore, and extended N-terminal region for Ca^2+^ mediated activation of this proteins is present only in plant Rboh proteins. However, the mammalian homologue catalytic unit gp91phox shares structural and functional domains with plant Rboh proteins (Cheng et al., 2013; Cheng et al., 2019). The hydrophilic C-terminal domain has cytosolic facing FAD and NADPH-binding sites. At the apoplast, heme groups are necessary for electron transport across the membrane to oxygen (O_2_, the electron acceptor) through flavin adenine dinucleotide (FAD) (Mahilingam et al., 2021).

Rboh of plants are a small multigene protein family (Marino et al., 2012). Till date, genes encoding *Rboh* or NADPH oxidase have been predicated and characterized in a number of plant species, namely *Citrus sinensis* (Zhang et al., 2022), *Capsicum annuum* (Zhang et al., 2021), *Hordeum vulgare* (Mahilingam et al., 2021, Lightfoot et al., 2008) *Nicotiana tobacum* (Yu et al., 2020), *Malus domestica*, *Prunus avium*, *Prunus dulcis*, *Rubus occidentalis*, *Fragaria vesca* and *Rosa chinensis* (Cheng et al., 2020), *Tritium aestivum* (Navathe et al., 2019 Hu et al., 2018), *Glycine* max (Liu et al., 2018), *Oryza Sativa* (Yamauchi et al., 2017, Wong et al., 2007) *Malus domestica* (Cepauskas et al., 2016) *Vitis vinifera* (Cheng et al., 2013), *Medicago truncatula* (Marino et al., 2011), *Zea mays* (Lin et al., 2009), *Arabidopsis thaliana* (Torres et al., <u>1998</u>; <u>2005</u>), *Lycopersicon esculentum* (Sagi and Fluhr, 2001). The rice OsRbohA was the first identified plant kingdom Rboh protein (Navathe et al., 2019). The genome of model plant *Arabidopsis thaliana* has ten *Rboh* genes, and as per GeneVestigator microarray datasets (Zimmermann et al., 2004), *AtRbohH* and *AtRbohJ* are involved with the growth of pollen tube tip, while *AtRbohA, AtRbohB, AtRbohC, AtRbohE, AtRbohG* and *AtRbohI* expressed in root tissues, and *AtRbohD* and *AtRbohF* across all the tissues (Hawamda et al., 2020). The member *AtRbohB* is involved in seed ripening and root hair formation, whereas *AtRbohC* controlled root hair cell growth (Navathe et al., 2019). Expression of *AtRbohD* and *AtRbohE* were also induced by jasmonic acid (JA) indicating their role in stress response and signalling (Maruta et al., 2011). Additionally, *AtRbohE* is also reported to regulate tapetal programmed cell death (PCD) and pollen formation in wheat (Hu et al., 2018).

The two species of the genus *Aquilaria*, viz. *A. agallocha* and *A. sinensis* (commonly known as agar), belong to the family Thymelaeaceae of the plant group Angiosperm (Kristanti et al., 2018). As a result of biotic and abiotic stress, the heartwood of *Aquilaria* tree transforms into worthy resinous dark wood known as agarwood (Das et al., 2021). Agarwood is popular globally for its use as raw material in perfume, incense, and medicine (Monggoot et al., 2018). When *Aquilaria* trees are physiologically triggered by physical wound, insect invasion or microbial infection, it starts to activate defence related signal transduction pathway to accumulate fragrant metabolites as a defence product (Tan et al., 2019; Liu et al. 2013; Mohamed et al., 2014). Sesquiterpenes and 2-(2-phenylethyl) and chromones are two prominent chemical constituents of agarwood which form occlusions of vessels, an efficient conducting structure of xylem and limits further infection (Naef et al., 2011; Chen et al., ^2^01^2^; Yang et al., 2021 Zhang et al., 2014). H_2_O_2_ burst (ROS production) is known to occur in wounded *Aquilaria* tree resulting in production of sesquiterpenes and other metabolites, and deposition of resins (Zhang et al., 2014). Also in other plants, H_2_O_2_ is known to play role in regulation during secondary metabolites viz., capsodiol (Tobacco), phenolics (Carrot), and β-coumaroyloctopamine (Potato) biosynthesis (Matsuda et al., 2001). Interestingly, in *A. sinensis*, upregulation of three sesquiterpene synthase, namely *AsTPS10*, *AsTPS16*, and *AsTPS19* leads to sesquiterpene accumulation in H_2_O_2_ pruned stem (Lv et al. in 2019). Additionally, methyl jasmonate could also upregulate expression of *Rboh* genes considerably in calli of *A. sinensis* (Xu et al., 2013). The Rboh enzymes has also been observed to play important role in the accumulation of 2-(2-phenylethyl) chromones (Wang et al., 2018). Various previous studies have established the role of *Rboh* gene families in ROS molecules accumulations after microbial attack in plants (Morales et al., 2016, Chang et al 2020, Pacheco Trejo et al., 2022). Since, agarwood is an outcome of microbes-mediated stress, it lead us to hypothesize that during microbial invasion, the *Aquilaria* plants accumulate ROS molecules through the action of Rboh proteins. The ROS produced initiate a cascade of biochemical reactions which activate the defence related signal transduction pathways which eventually activate genes involved in biosynthesis of secondary metabolites for defense responses (Xu et al., 2016, Tan et al., 2019, Das et al., 2021,).

To best of our knowledge, study of Rboh protein family related to defensive mechanism in *Aquilaria* has not been reported. Therefore, systemic identification and classification of *Rboh* members in genome-wide basis, and their expression analysis in stress induced tissue will likely identify the key members involvement in ROS generation and stress response in *Aquilaria*. Additionally, the findings in this study will also lay the groundwork for understanding the molecular basis and regulatory mechanisms of *Rboh* genes and their possible involvement in agarwood deposition in *Aquilaria* species.

## 2. Materials and methods

### 2.1. Sequence Retrieval and Identification of the *Rboh* genes

The predicted protein sequences of *A. agallocha* and *A. sinensis* were obtained from previous annotation projects (Das et al., 2021 and Nong et al., 2020). The respiratory burst NADPH oxidase domain (PF08414), ferric reductase NAD binding domain (PF08030), FAD-binding domain (PF08022), and ferric reductase like transmembrane component (PF01794) were downloaded from the Pfam database to build Hidden Markov Models (HMM) profile with HMMER 3.3.2. To identify *AaRboh* and *AsRboh* genes, protein sequences were searched against the build HMM model with hmmsearch (Potter et al., 2018). Thereafter, the redundant protein sequences were discarded. All the putative sequences were further confirmed through Pfam database (available online: http://pfam.xfam.org/) and SMART database (available online: http://smart.embl-heidelberg.de/). The physiological properties such as molecular weight (kDa), isoelectric point (pI), instability index (II), aliphatic index, and the grand average of hydropathicity (GRAVY) were calculated by using the ExPASy-ProtParam tool (available online: http://web.expasy.org/protparam/).

### 2.2. Multiple sequence alignments and phylogenetic analysis

Rboh proteins of *Solanum tuberosum*, *Arabidopsis thaliana*, *Hordeum vulgare,* and *Glycine max* were downloaded from NCBI and aligned in MEGA-X program with predicted AaRboh and AsRboh protein sequences. Phylogenetic tree was constructed in MEGA-X (available online at https://www.megasoftware.net/) program using neighbour joining method based on the P distance model with a bootstrap of 1000 replicates (Kumar et al., 2016). For the conserved domain analysis sequences were aligned using Clustal X and presented with ESPrit 2.2-ENDscript 1.0 (Robert et al., 2014).

### 2.3. Subcellular location, regulatory elements, Gene structure, and Conserve motif

The subcellular location of all retrieved Rboh proteins was predicted with online web server CELLO version 2.5 (http://cello.life.nctu.edu.tw/). The intron-exon structure of individual *Rboh* genes were predicted using the corresponding genomic DNA sequences and CDS sequences as input to Gene structure Display Server GSDS2.0 (available online http://gsds.cbi.pku.edu.cn) (Hu et al., 2015). The conserved motifs in *Rboh* genes were predicted by the MEME suite (http://meme-suite.org/) with the default parameters (Bailey et al., 2009). The cis-acting regulatory elements were searched in the 2400 bp upstream of each gene start site in the two *Aquilaria* genomes using the PLANT CARE database (Lescot et al., 2002).

### 2.4. Functional predictions and Protein-protein interactions

Probable functions of the AaRboh proteins were predicted based on the GO terms and KEGG annotations and their functional partners were analysed through STRING v11.5 program with default parameters utilizing *Arabidopsis* homologous proteins.

### 2.5. Synteny analyses and Ka/Ks calculation

Synteny blocks within the two *Aquilaria* genomes, and with other plant (*A. thaliana* and *S. tuberosum*) were constructed using Quick MCScanX Wrapper and were visualized using the Dual Synteny plotter in TBtools (Chen et al., 2020). Gene duplication events were predicted by the blastP program with the parameters **≥** 80% of sequence identity and alignment coverage. Homologous genes pair within 100 kb region in the same chromosome designated as tandem duplication, while genes located beyond 100 kb as segmental duplication (Islam et al., 2019). The non-synonymous substitution rate (Ka), synonymous substitution rate (Ks), and Ka/Ks ratio were calculated with TB tools (Wang et al., 2010). Divergence time for duplication of paralogs pairs of the gene was calculated following the formula T = Ks/2λ (where λ indicates the clock-of-like rate of 6.96 synonymous substitutions per 10^−9^ years (Lopez-Ortiz et al., 2019).

### 2.6. Expression patterns of the *AaRboh* and *AsRboh* genes

To study transcript abundance of *Rboh* genes, the RNA-Seq data was extracted from the NCBI Geo website (https://www.ncbi.nlm.nih.gov/sra) and short reads were aligned to the genome using HISAT2 (Kim et al., 2015), and then the reads were assembled and quantified using StringTie (Kovaka et al. 2019). Expression values for individual *Rboh* genes were calculated as fragments per kilobase of exon model per million mapped reads (FPKM). The transcript abundance differences of *AaRboh* genes between agarwood (SRX4149019-SRX4149021) and healthy (SRX4184708-SRX4184710) wood tissues of *A. agallocha* was also calculated. Similarly, transcript abundance of *AsRboh* genes were determined in the different tissues/organs viz., aril (SRX6871071, SRX6871068), seed (SRX6871057, SRX6871070), flower (SRX6871060, SRX6871063), bud (SRX6871059, SRX6871062), leaf (SRX6871066, SRX6871058), salinity stressed callus (SRX1495981, SRX1495736), flower (SRX6871059, SRX6871062) and wounded stem (SRX6871056, SRX6871064) of *A*. *sinensis*.

### 2.7. Plant material, growth and treatment

*A. agallocha* calli were induced from leaves on Murashige-Skoog (MS) medium containing 6 mg/L dichlorophenoxyacetic acid (2,4-D) and 2 mg/L kinetin (KT). The calli were transferred to fresh MS medium after every month till formation of friable calli. In order to induce stress, the calli were transferred to an MS medium containing 10 mM H_2_O_2_ and was also exposed to 5 mM dimethylthiourea (DMTU, an H_2_O_2_ scavenger) (Wang et al., 2018), and calli without any treatment served as control. The samples were harvested at 0, 1, 2, 6, 12, 24, and 48 h post treatment.

Healthy saplings of *A. agallocha* maintained in pots at Bioengineering and Technology department of Gauhati University were selected for stress treatments as per standard methodology (Lv et al., 2019). The lateral stems were cut with scissors, and one cm from the apical end of cut lateral stems was immersed separately in distilled H_2_O, H_2_O_2_, DMTU solutions. Thereafter, the pre-treated 1 cm portions were discarded and on the top of the treated stems (approximately 2 cm) were exposed to air. Samples were harvested after 0, 1, 2, 6, 12, 24, and 48 h of air exposure. Treatment of seedlings after cutting refers to physical wounding. For the stress treatment of seedlings, the wounded stems were treated with distilled H_2_O, H_2_O_2_, DMTU and healthy *Aquilaria* lateral stems were taken as a control. For analysis of *AaRboh* gene transcript in resin embedded wood tissue (induced) and healthy wood tissue of *Aquilaria* tree were collected from the Hoollongapar Gibbon Sanctuary in Jorhat, Assam, India, using methodology outlined previously (Islam et al 2020). All sample sets were rapidly immersed in liquid nitrogen and stored at 80 °C for further experiments.

### 2.8. RNA extraction and qRT-PCR analysis

Total RNA was extracted from stem tissues using the RNA extractions method as described by Islam and Banu, 2019 and from callus tissues using RNeasy Plant Mini Kit (Qiagen).The quality and concentration of extracted RNA were checked with 1 % agarose gel electrophoresis and estimated with Multiskan Sky Microplate Spectrophotometer (Thermo Fisher Scientific, USA), respectively. A total of 1 µg of RNA was used to synthesize first strand cDNA with SuperScript III Reverse Transcriptase (Thermofisher). qRT-PCR was performed using QuantStudio™ 3 real-time PCR system (Applied Biosystems, USA) and PowerUp SYBR Green Master Mix of Applied Biosystems. Gene specific primers for *AaRboh*s were designed with PrimerQuest software of IDT (https://sg.idtdna.com/pages/tools/primerquest/) and are enlisted at Supplementary file 1. The standardized GAPDH primer of *A. malaccensis* was used as internal control (Islam et al., 2020). For each biological replicate, the analyses were performed with three technical replicates each containing 20 µl of reaction volume in optical stripes, the temperature pattern of 95 °C for 1 min, followed by 40 cycles at 95 °C for 10 s and 60 °C for 30 s was followed as thermal cycler profile. Fold change in the gene expression was determined by ΔΔCt method.

### 2.9. ROS determination of treated plant materials

The endogenous ROS production was determined according to Wang et al., 2018 with slight modifications. Plant samples (3 g of fresh weight) were subjected to individual and combined treatments with H_2_O_2_ and DMTU. The treated samples were homogenized in 3 mL of pre-cooled acetone using mortar and pestle on ice. Obtained mixtures were centrifuged at 3000 rpm for 10 min at 4 °C. The supernatant obtained (0.1 mL) was quickly mixed with 0.1 mL of 5 % TiSO_4_ and to it 0.2 mL of NH_4_OH_2_ was added. The resulting mixture was centrifuged at 3000 rpm at 4 °C for 10 min to separate the titanium-hydroperoxide complex precipitate and supernatant was discarded. After three washes with pre-cooled acetone, the precipitate was dissolved in 2 mL of 2 mol/L of H_2_SO_4_. The absorbance of the dissolved precipitate at 415 nm was monitored to quantify H_2_O_2_ content. Obtained absorbances were compared with calibration curve derived from known concentrations of freshly prepared 30% H_2_O_2_ (Wang et al 2018, Zhang et al., 2021).

## 3. Results

### 3.1. Identification of the *Rboh* genes in *Aquilaria* species

Fourteen *Rboh*s genes were identified from the two *Aquilaria* genomes, 7 each from *A. agallocha (AaRboh)* and *A. sinensis (AsRboh)* (Supplimentary file 2). *Aquilaria* species have a total 2n =16 number of chromosomes and *AsRbohs* were found to be distributed on five chromosomes (Chr02, Chr04, Chr06, ContigUN and Chr07), and *AaRbohs* on seven different scaffolds (KK901300.1, KK899295.1, KK902390.1, KK900302.1, KK899913.1,

KK900079.1, and KK899008.1) as chromosome level assembly was missing in *A. agallocha*. Amongst all 14 predicted proteins, AaRbohJ had the shortest protein sequence (663 amino acids), whereas AaRbohA had longest protein (946 amino acids) (Table. 1). The molecular weights and pIs ranged from 75.72 kDa (AaRbohJ) to 107.77 kDa (AsRbohA) and from 8.93 kDa (AsRbohC2) to 9.45 kDa (AaRbohA). Based on the instability index (II) the 3 proteins (AsRbohC1, AsRbohC2, and AaRbohC) could be categorised as stable, and the rest 11 as thermostable, according to the aliphatic index (Ai). Grand average of hydropathicity (GRAVY) values of all predicted Rboh proteins were less than zero, indicating that they are hydrophilic in nature.

**Table 1:**
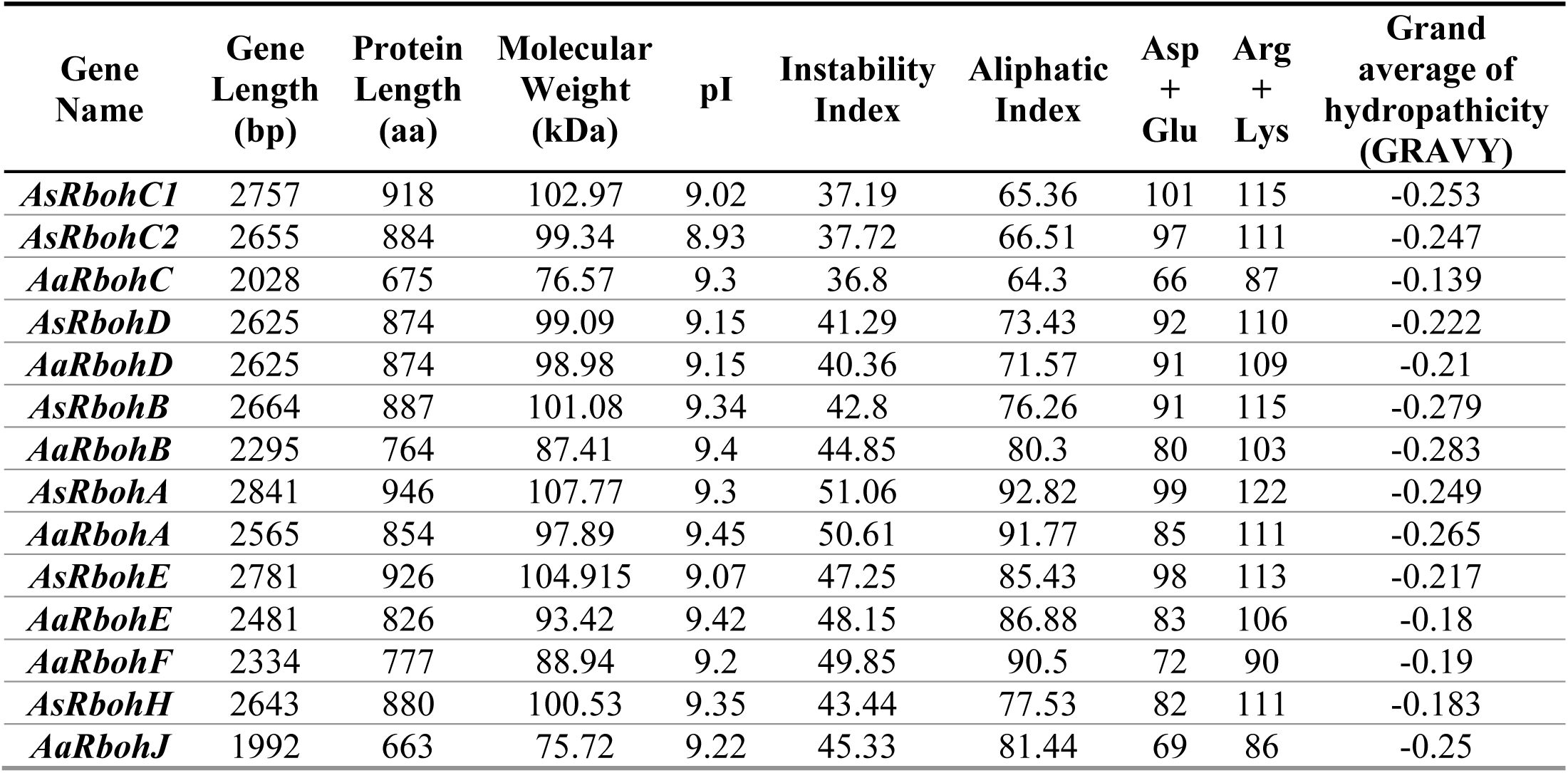
Details of the 14 putative *Rboh* genes identified in this study

| Gene Name | Gene Length (bp) | Protein Length (aa) | Molecular Weight (kDa) | pI | Instability Index | Aliphatic Index | Asp + Glu | Arg + Lys | Grand average of hydropathicity (GRAVY) |
| --- | --- | --- | --- | --- | --- | --- | --- | --- | --- |
| *AsRbohC1* | 2757 | 918 | 102.97 | 9.02 | 37.19 | 65.36 | 101 | 115 | -0.253 |
| *AsRbohC2* | 2655 | 884 | 99.34 | 8.93 | 37.72 | 66.51 | 97 | 111 | -0.247 |
| *AaRbohC* | 2028 | 675 | 76.57 | 9.3 | 36.8 | 64.3 | 66 | 87 | -0.139 |
| *AsRbohD* | 2625 | 874 | 99.09 | 9.15 | 41.29 | 73.43 | 92 | 110 | -0.222 |
| *AaRbohD* | 2625 | 874 | 98.98 | 9.15 | 40.36 | 71.57 | 91 | 109 | -0.21 |
| *AsRbohB* | 2664 | 887 | 101.08 | 9.34 | 42.8 | 76.26 | 91 | 115 | -0.279 |
| *AaRbohB* | 2295 | 764 | 87.41 | 9.4 | 44.85 | 80.3 | 80 | 103 | -0.283 |
| *AsRbohA* | 2841 | 946 | 107.77 | 9.3 | 51.06 | 92.82 | 99 | 122 | -0.249 |
| *AaRbohA* | 2565 | 854 | 97.89 | 9.45 | 50.61 | 91.77 | 85 | 111 | -0.265 |
| *AsRbohE* | 2781 | 926 | 104.915 | 9.07 | 47.25 | 85.43 | 98 | 113 | -0.217 |
| *AaRbohE* | 2481 | 826 | 93.42 | 9.42 | 48.15 | 86.88 | 83 | 106 | -0.18 |
| *AaRbohF* | 2334 | 777 | 88.94 | 9.2 | 49.85 | 90.5 | 72 | 90 | -0.19 |
| *AsRbohH* | 2643 | 880 | 100.53 | 9.35 | 43.44 | 77.53 | 82 | 111 | -0.183 |
| *AaRbohJ* | 1992 | 663 | 75.72 | 9.22 | 45.33 | 81.44 | 69 | 86 | -0.25 |

### 3.2. Multiple Sequence Alignment and Phylogenetic analysis

To identify the phylogenetic positions of the identified *Aquilaria* members, a tree was constructed for 7 Rboh proteins each of *A. agallocha* and *A. sinensis,* and 5, 12, 16, 16 proteins of *S. tuberosum*, *A. thaliana*, *H. vulgare* and *Glycine max,* respectively. The analysis showed clustering of members of the six-plant species into five main groups (Fig. 1). The highest numbers of members were distributed in Group 1(18) and Group 3 (15), followed by Group 2 (13), Group 4 (9), and Group 5 (8). Distribution of *Aquilaria* Rboh proteins based on their phylogenetic positions in these groups were as follows: Group 1 comprised of 2 AaRboh (AaRbohD, AaRbohC); 3 AsRboh (AsRbohD, AsRbohC1-C2) protein; 2 StRboh (StRbohC, StRbohD); 4 GmRboh (GmRbohC1-C4); 4 AtRboh (AtRbohA, AtRbohC, AtRbohD, AtRbohG); 3 HvRboh (HvRbohI1-I3). While Group 2 comprised of 5 HvRboh (HvRbohB1-B4, HvRbohH); 4 GmRboh (GmRbohB1-B4); 1 each of AaRboh (AaRbohB), AsRboh (AsRbohB), AtRboh (AtRbohB), StRboh (StRbohB). Similarly, Group 3 comprised of 4 each of HvRboh (HvRbohF1-F2, HvRbohA1-A2) and AtRboh (AtRbohF1-F3, AtRbohI); 3 GmRboh (GmRbohA1, GmbohA3-4); 2 StRboh (StRbohA, StRbohF); 1 each of AaRboh (AaRbohA) and AsRboh (AsRbohA). Group 4 comprised of 4 GmRboh (GmRbohE1-E2, GmRbohF1-F2); 2 AaRboh (AaRbohE, AaRbohF); and 1 each of AsRboh (AsRbohE), AtRboh (AtRbohE) and HvRboh (HvRbohE2). Group 5 comprised of 3 HvRboh (HvRbohD, HvRbohJ, HvRbohE1); 2 AtRboh (AtRbohH, AtRbohJ); 1 each of AaRboh (AaRbohJ), AsRboh (AaRbohH), and GmRboh (GmRbohH1). Al the 5 groups comprised of atleast one members of 6 the plants including *A. agallocha* and *A. sinensis* except *S. tuberosum* where no member belonged to Group 4 and 5.

**Fig 1.**
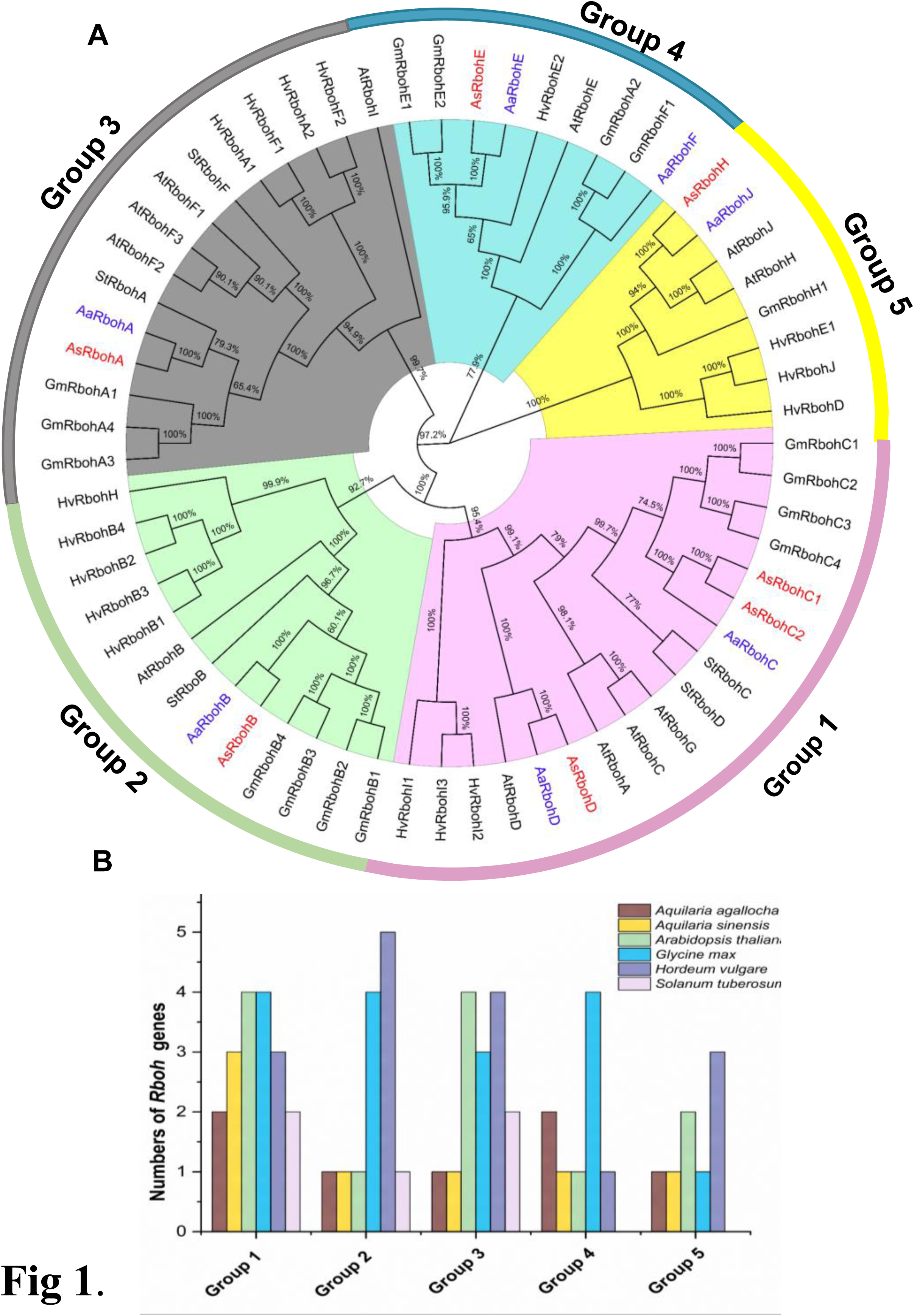
Phylogenetic relationship (A) Molecular phylogenetic tree of Rboh proteins of *Aquilaria agallocha, Aquilaria sinensis, Arabidopsis thaliana, Glycine max, Hordeum vulgare, Solanum tuberosum* was constructed using MEGA-X with NJ methods-based P distance substitutions model. Bootstraps values used to assess the tree. Five group are shown as Group1-5 with different colour. (B) The numbers of Rboh protein sequences in the Group 1-5.

### 3.3. Subcellular Location, Gene structure, and Conserve motif

Subcellular location prediction showed that all the Rboh protein are located on the plasma membrane. The genes clustered together tend to possess a similar organization of exon-intron structures and indicates close evolutionary relation. The coding sequences (CDS) of putative *Rbohs* had 8 (*AaRbohD*, *AsRbohD*) to 15 (*AaRbohE*) numbers of exons whereas most of the predicted *Rboh* genes contained 12 (*AaRbohB*, *AaRbohA*, *AsRbohC1*) or 14 (*AaRbohF AsRbohB*, *AsRbohC2*, *AsRbohA*, *AsRbohE*) number of exons (Fig. 2). The number and types introns varied in the members. Highest was observed for phase 0 introns (81), followed by phase 2 introns (42), and phase 1 introns (31). The number of phase 0 introns varied from two to nine in each member, while phase 1 introns from one to three, and phase 2 introns from two to three phase 2 introns (except *AsRbohJ*) (Fig. 2).

**Fig 2.**
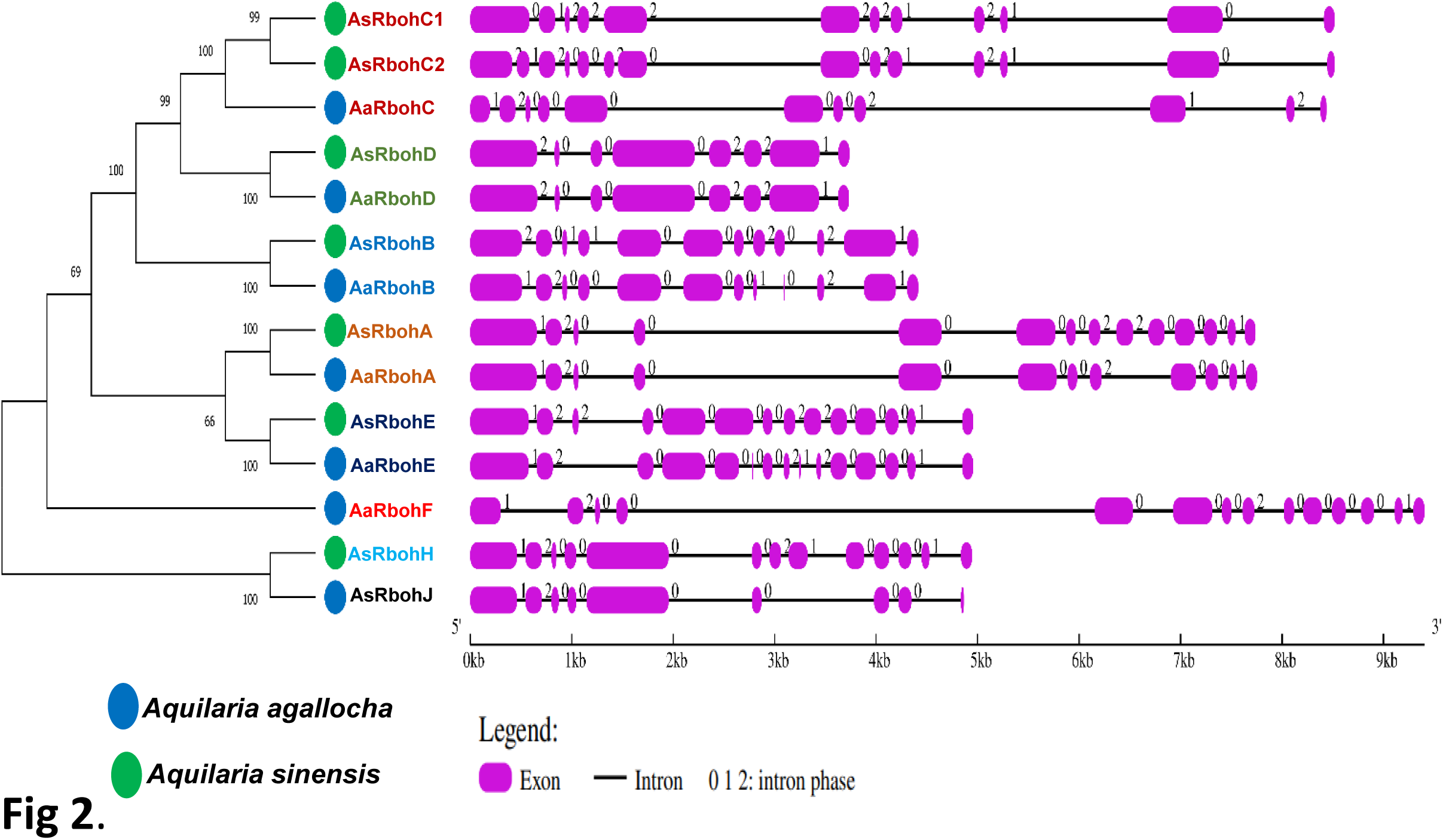
Schematic representation of structures of 14 putative *Rboh* genes in two *Aquilaria* species. The exons and introns indicated with pink rectangles and black colour lines respectively on the right of phylogenetic tree. The numbers (0, 1 and 2) given to the gene’s structures indicate their respective intron phases. The sizes of exon-intron can be directed as indicated in scale at the bottom.

### 3.4. Amino Acid Sequence and Characteristic Domain Analysis of *Rboh*s

Ten consensus motifs were identified through MEME suite in the Rboh proteins based on conserved amino acid residues. The motifs viz. motif 1 (except AaRbohJ), motif 2, motif-3 (except AaRbohE), motif 4, motif 5, motif 6, motif 7, and motif 8 existed in all Rboh proteins (Fig. 3A). Whereas, both motif 9 and 10 were missing in *Aquilaria* RbohB, RbohC, and RbohJ. Interestingly, the members of Group 1 (AsRbohC1, AsRbohC2, AaRbohD, and AsRbohD); Group 2 (AsRbohB); Group 3 (AsRbohA); Group 4 (AsRbohE, AaRbohF); and Group 5 (AsRbohH) contained the highest i.e., all the ten motifs. The four conserved motifs typically found in Rboh proteins which are required for catalysis were also existed in the *Aquilaria* Rboh members (Fig. 3B). The most conserved amino acid within these motifs were also identified which were represented by higher bits size (Fig. 3C). Alignment of the amino acid sequences of the *Rboh* homologs of *A. agallocha* and *A. sinensis* with Clustal W and ESPrit 2.2-ENDscript 1.0 revealed the presence of conserved domains i.e., NADPH oxidase (PF08414), EF hand, Ferric reductase (PF01794), FAD binding (PF08022), and NAD binding (PF08030) in all the 14 Rboh protein sequences (Fig. 4). However, NAD binding domain was missing in AaRbohB, AaRbohF, and AaRbohJ, and Ca^2+^-binding EF-hand domain in AaRbohF.

**Fig 3.**
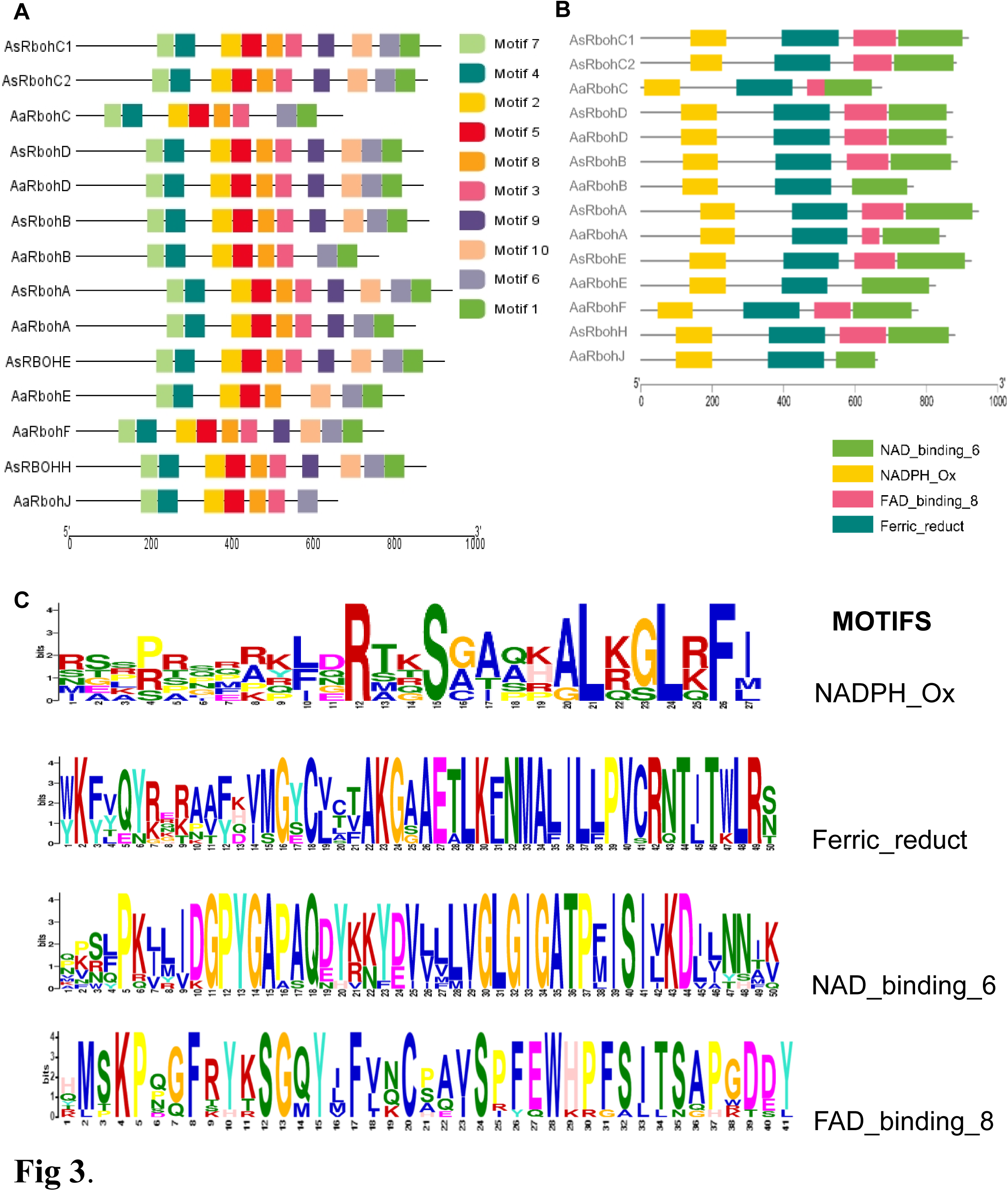
Conserved motifs distribution (A) Ten types of conserved motifs of *AsRboh* and *AaRboh*. (B) four particular characteristics motif of *AsRboh* and *AaRboh*. (C) Sequence logos of the NADPH_Ox, Ferric_reduct, FAD_binding_8, and NAD_binding_6 conserved motifs in *Rboh* proteins are shown in (a), (b), (c), and (d), respectively. The bit score identifies the amount of information present at each position in the sequence. The degree of amino acid conservation across all proteins tested is correlated with the height of each character.

**Fig 4.**
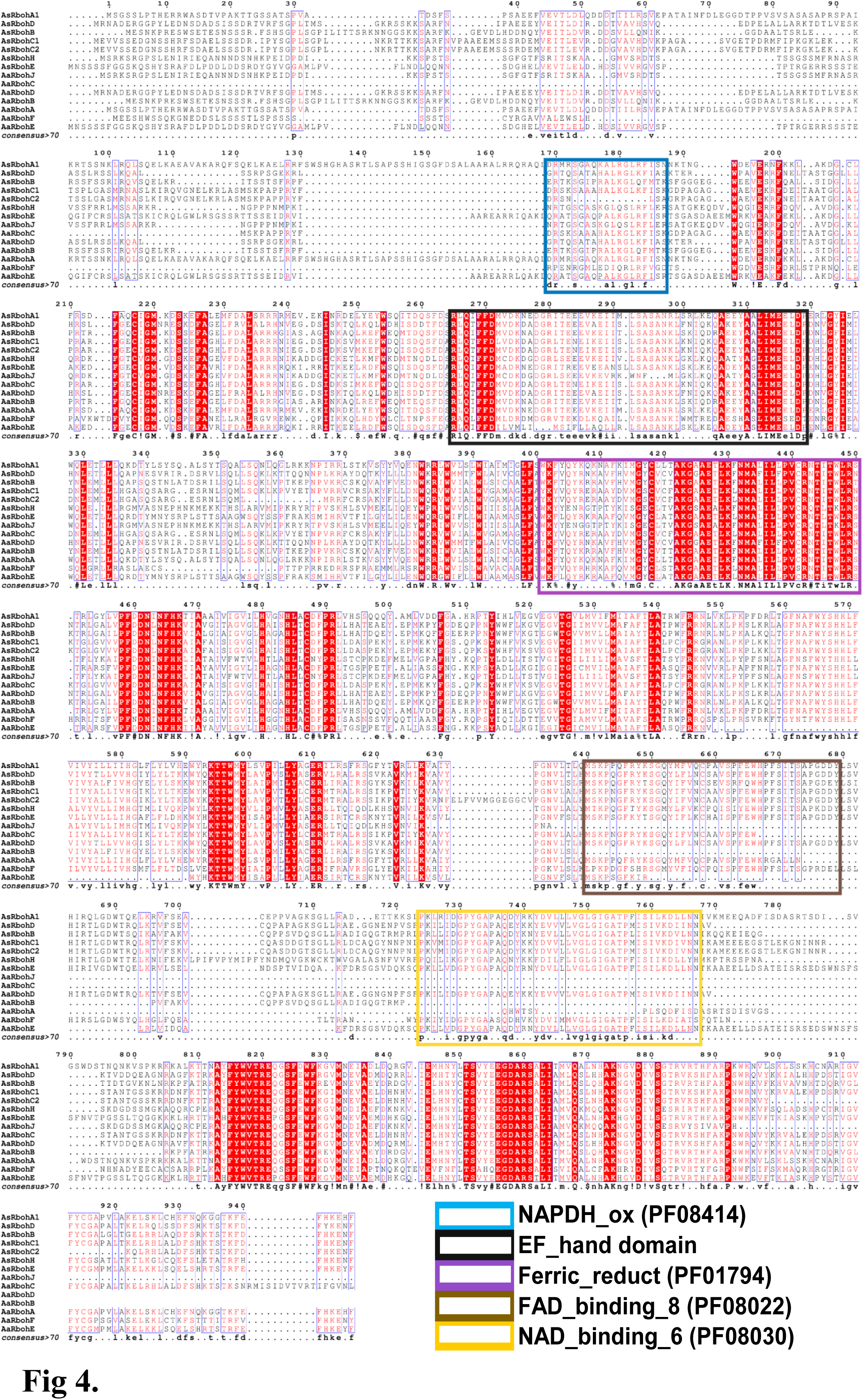
Multiple protein sequence alignment and domain structure of Rboh proteins of *A. agallocha* and *A. sinensis*. Highly conserved amino acids indicate with red shading and low level of amino acids represents with lighter shading. The NADPH_ox (PF08414), EF-hand domain, Ferric_reduct (PF01794), FAD_binding_8(PF08022) and NAD_binding_6 (PF08030) indicates with blue, black, violet, brown and yellow colour, respectively.

### 3.5. Cis-Acting Elements in the putative *Rboh* promoter

A total of 56 types of cis-acting elements were identified in the 2.4 kb region upstream of the translational start site of each *Rboh* genes (Supplementary file. 3). These elements could be categorised into four major functional groups: stress response, hormone regulation, and development. Defense and stress responsiveness cis-acting element were ARE (cis-acting elements for anaerobiosis), MBS (drought response), LTR (low-temperature responsive cis-acting element), TC-rich repeats (defence), and WUN motif (wound stress) (Fig. 5A). Additionally, 6 types of plant hormone regulatory elements, namely salicylic acid response element (TCA-element and SARE); Gibberellin response element (TATC-box, P-box, and GARE); auxin response element (AuxRR-core, TGA-element); ethylene response element (ERE); abscisic acid (ABA) response element (ABRE); and methyl jasmonate response elements (TGACG-motif, CGTAC-motif) were also found in the putative *Rboh* gene promoters (Fig. 5A). The promoters also had RY-elements, CAT box cis-elements which were associated with cell differentiation and developmental processes. The numbers defense and stress responsiveness elements varied from 6 (*AaRbohE*) to 14 (*AaRbohA*) and were observed in all the members (Fig. 5B). Most of the members had methyl jasmonate and other hormone response elements indicating their inducible nature.

**Fig 5.**
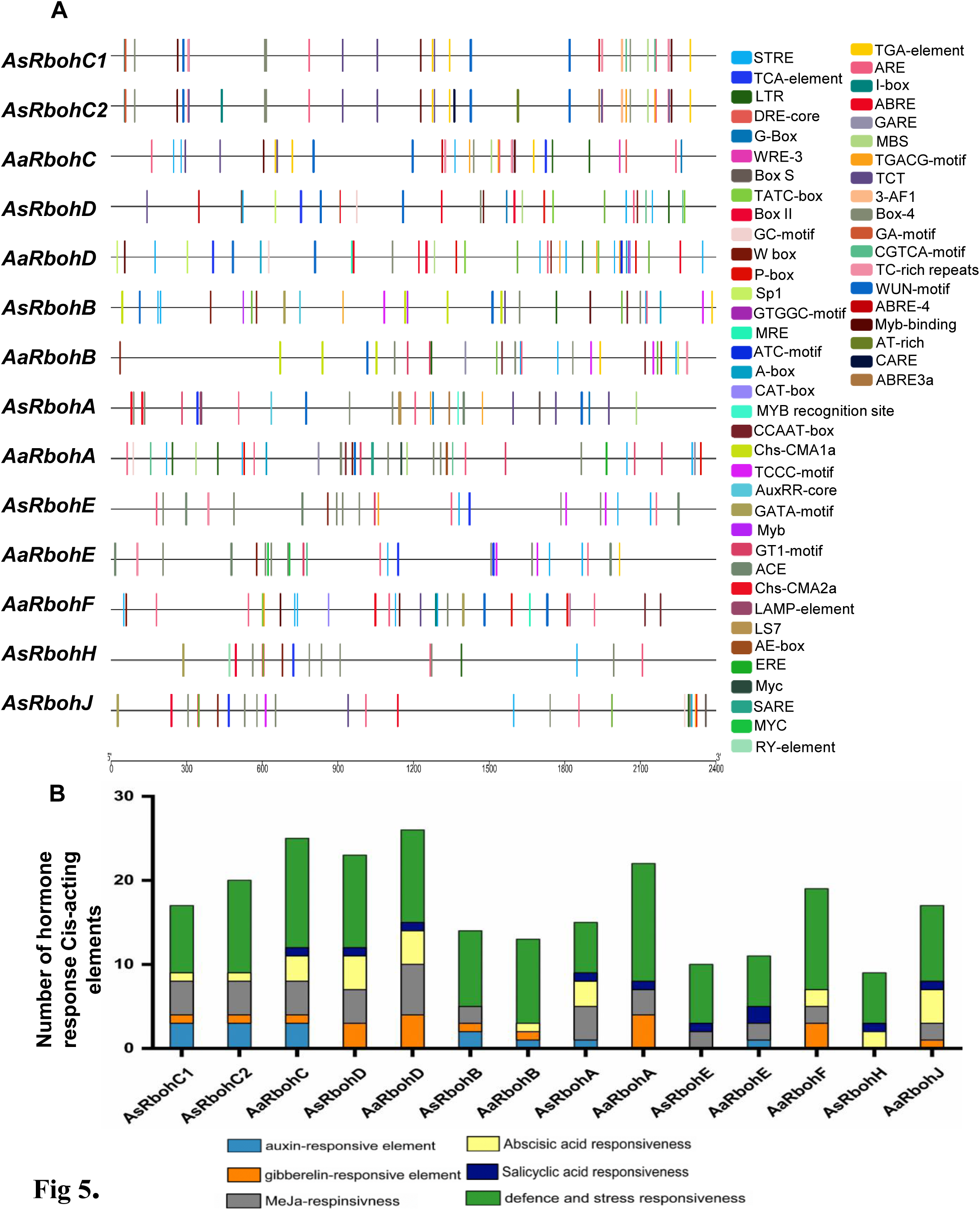
Cis acting regulatory elements (A) Total cis-acting regulatory elements prediction in the 2.4 kb putative promoters of *AaRboh* and *AsRboh* genes. different coloured rectangles lines indicate the different types of cis-acting elements of biotic, abiotic, growth and developments (B) cis-acting regulatory elements responsible for different types of hormonal responses, defence and stresses. cis-acting elements with same or similar functions represents with same colour.

### 3.6. Gene location, synteny analysis and Ka/Ks calculation

The seven *Rboh* gene of *A. agallocha* were found to be distributed in seven different scaffolds (Fig. 6A), while in *A. sinensis*, as chromosome level assembly is available, were found to be distributed in Chromosome 2 (*AsRbohA*), Chromosome 4 (*AsRbohE*), Chromosome 6 (*AsRbohB*), Chromosome 7 (*AsRbohC1*, *AsRbohD*), and ContigUN (*AsRbohC2*) (Fig. 6B). The syntenic study revealed that the five *AaRboh* genes viz. *AaRbohA, AaRbohB, AaRbohC, AaRbohF,* and *AaRbohE* in *A. agallocha* had a collinear relationship with members of *A. sinensis*. Interestingly, *AaRbohE* was associated with two syntenic gene pairs with *A. sinensis* (Fig. 6C). However, the *AaRbohD,* and *AaRbohJ* genes had no collinear genes with *A. sinensis*. Additionally, *AaRbohA* and *AaRbohC* showed collinear relationship with *A. thaliana*, and *AaRbohE* with *S. tuberosum*. The member *AsRbohC2* showed synteny with genes of both *S. tuberosum* and *A. thaliana*. The syntenic gene pairs with their genome location were summarized in Supplementary file Table 5. Duplication analysis showed that the pairs *AsRbohC1* and *AsRbohC2* underwent segmental duplication over the course of evolution. The Ka/Ks ratio of the duplicated gene pair was found to be < 1 (Supplementary file Table 4) indicating purifying selection during duplication.

**Fig 6.**
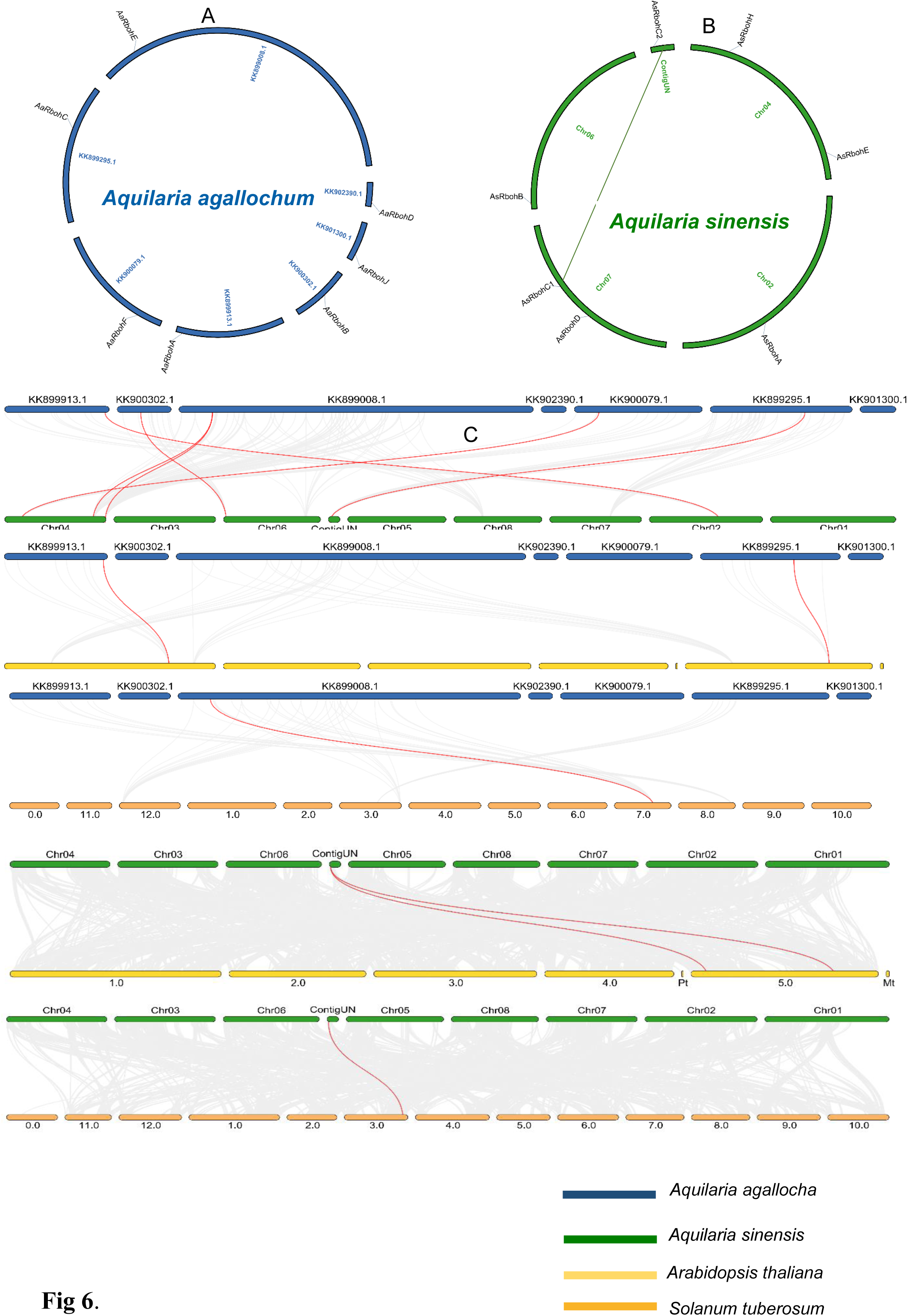
Synteny analysis (A) Synteny analysis of *AaRboh* (B) Synteny analysis of *AsRboh* and green line shows duplicate genes of *AsRboh* (C) Synteny analysis among *A. agallocha* and *A. sinensis; A. agallocha* and *A. thaliana; A. agallocha* and *S. tuberosum; A. sinensis* and *A. thaliana; A. sinensis and S. tuberosum.* Grey lines represent all collinearity blocks whereas red lines show orthologous gene pairs among two species.

### 3.7. In-silico expression *Rboh* genes in RNA-seq data of different tissues

Analyses of expression of Rboh genes in RNA-seq data of healthy and agarwood revealed *AaRbohC* expression to be the highest in agarwood tissue followed by *AaRbohA* (Fig. 7A). In contrast, *AaRbohE* and *AaRbohF* genes showed high expression in healthy tissues, while *AaRbohB* and *AaRbohE* expressed in both healthy and agarwood tissue. However, negligible expression of *AaRbohD* and *AaRbohJ* was observed in both tissues. In addition, transcript abundance of *AsRboh* in RNA-seq data of different tissues including aril, seed, flower bud, leaf, callus, flower and wounded stem was estimated in order to identify tissue specific expression of the members. *AsRbohB* was found to express specifically in salt treated callus and slightly in wounded stem, and not in any other tissue (Fig. 7B). Similarly, *AsRbohD* also expressed highly in wounded stem, followed by callus and flower. *AsRbohE* showed little expression in leaf, flower and wounded stem. The genes *AsRbohA* and *AsRbohC2* showed highest expression in callus tissues followed wounded stem, both these genes expressed negligibly in flower, flower bud, leaf, seed and aril. The *AsRbohC1* showed highest expression in wounded stem, followed by callus, flower, flower bud, seed aril and leaf. To be noted, both the paralogs *AsRbohC1* and *AsRbohC2* more or less expressed in all tissues except in leaf. However, *AsRbohH* showed no expression in any of the tissues. The expression values of all the genes in the different tissues were summarized in Supplementary file 6).

**Fig 7.**
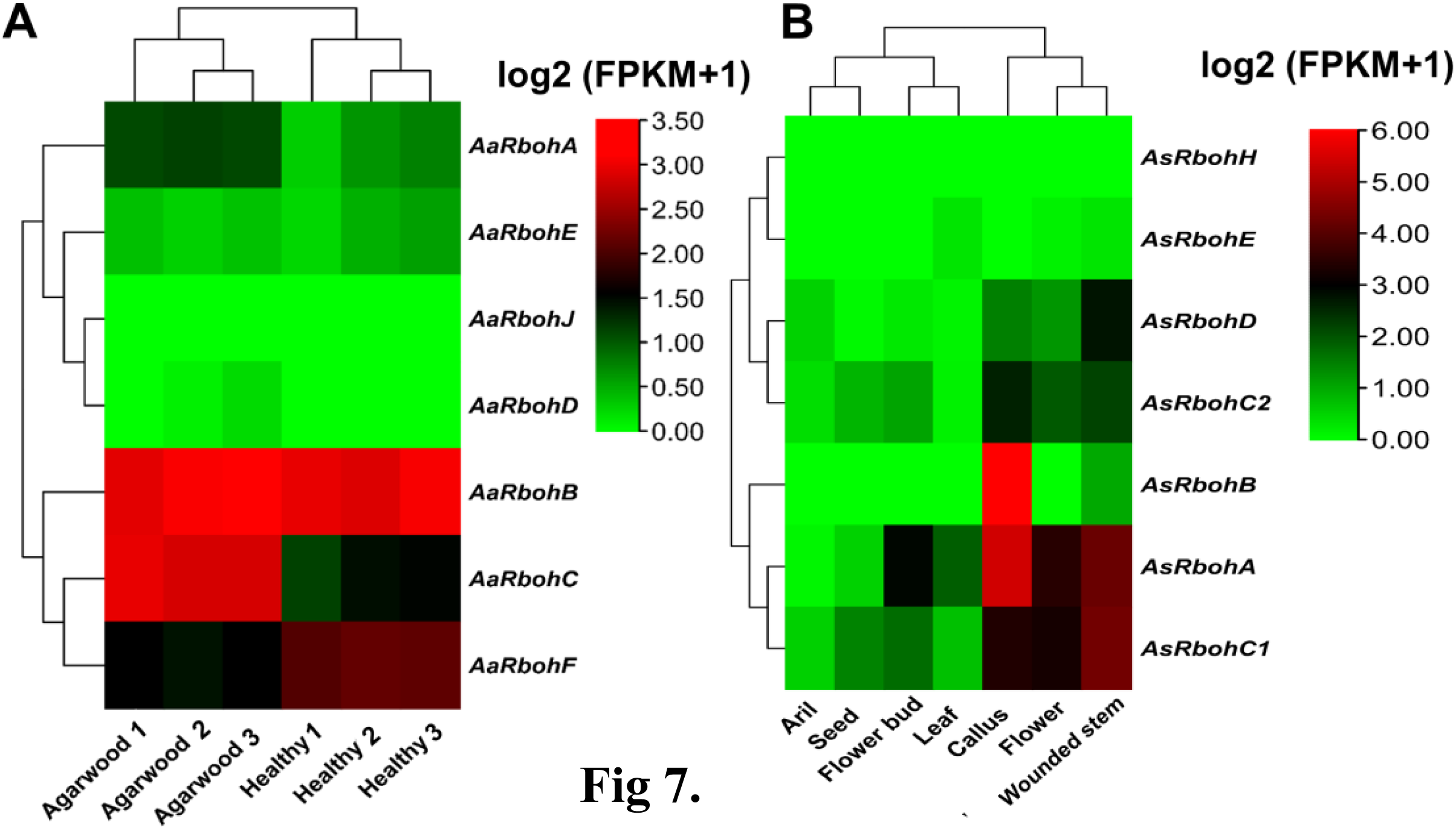
Expression profile (A) *AaRboh* gene expression patterns in healthy tissues and agarwood tissues are shown on a heat map. (B) *AsRboh* gene expression patterns in the 7 types tissues. All gene expression levels were converted to log scores value ranging from 0 to 6, and the corresponding low, moderate, or high expression levels were represented by green, black or red, respectively.

### 3.8. Transcript abundance of *AaRboh* genes upon H_2_O_2_ treatment

In order to investigate the effect of H_2_O_2_ in the expression of *AaRboh* genes, callus and wounded stem tissues were treated with H_2_O_2_, DMTU, combination of both, and H_2_O. In callus, it was observed that upon H_2_O_2_ exposure, expression of *AaRbohA* and *AaRbohC* peaked at 2 h (4.18-fold) and 6 h (24.40-fold), respectively. However, there was a steady decline in the expression for both the genes from 6 h to 48 h (Fig. 8). Furthermore, treatment with the combination of H_2_O_2_ and DMTU reduced their expression, and when treated with DMTU alone, expression was even lower than the control. The gene *AaRbohB* also showed similar expression pattern under the treatment condition. Interestingly, no difference in the expression pattern was observed in the three genes i.e., *AaRbohD, AaRBohF,* and A*aRbohJ* to that of the control throughout the time period. In wounded stem, the abundance of *AaRbohC* raised to 6.05-fold at 1 hr and subsequently returned to its initial level after 2 h (Fig. 9). After 6 h, its expression again increased to 6.6 times (40.21-fold change), but declined after 24 h of air exposure, while, there was no change in its expression in the healthy (control) stems. The *AaRbohA* also showed a similar trend of expression pattern, and to be noted, *AaRbohC* and *AaRbohA* elevated upon H_2_O_2_ treatment at 6 h and remained elevated even after 12 h, whereas DMTU treatment inhibited them back to 5-10-fold (Fig. 9). However, expression level was negligible in the healthy stems. Besides, *AaRbohB*, *AaRbohD*, *AaRbohF* and *AaRbohJ* showed no difference in the expression’s levels to that of the control.

**Fig 8.**
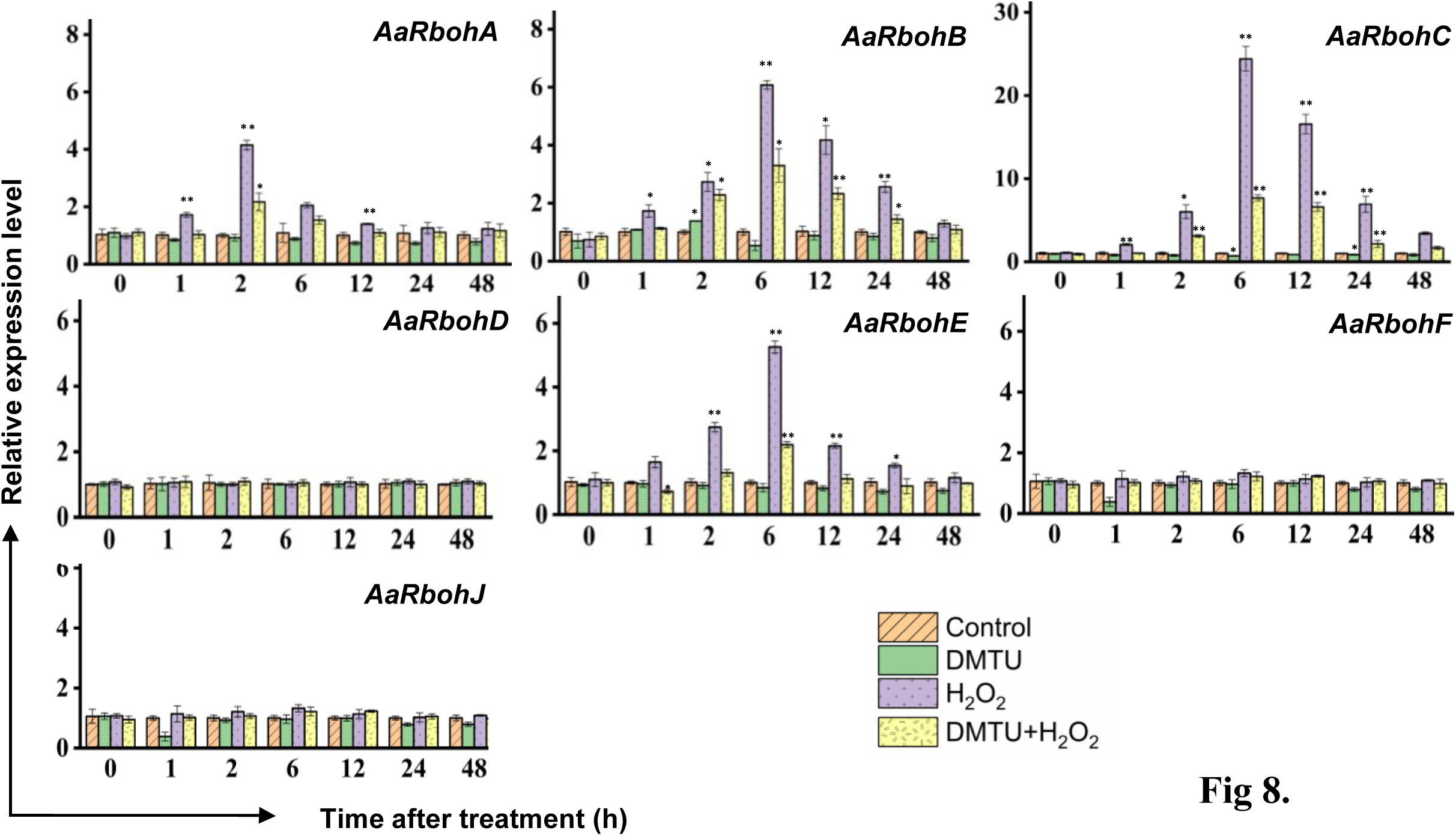
Relative expression levels of *AaRboh* genes in treated calli. relative transcripts abundance of 7 *AaRboh* were measured in calli tissue transferred to MS media with H_2_O_2_, H_2_O_2_+ DMTU, DMTU, respectively and calli without any treatment considered as control condition with samples harvested at 0, 1, 2, 6, 12, 24 and 48 h. Transcript abundances were measured using *A. agallocha* GAPDH as internal control. Asterisks (*) denotes a significant difference compared with healthy samples at 0.05 or **P < 0.01 (Student’s t-test). Data represent means ± SE off three independent experiments.

**Fig 9.**
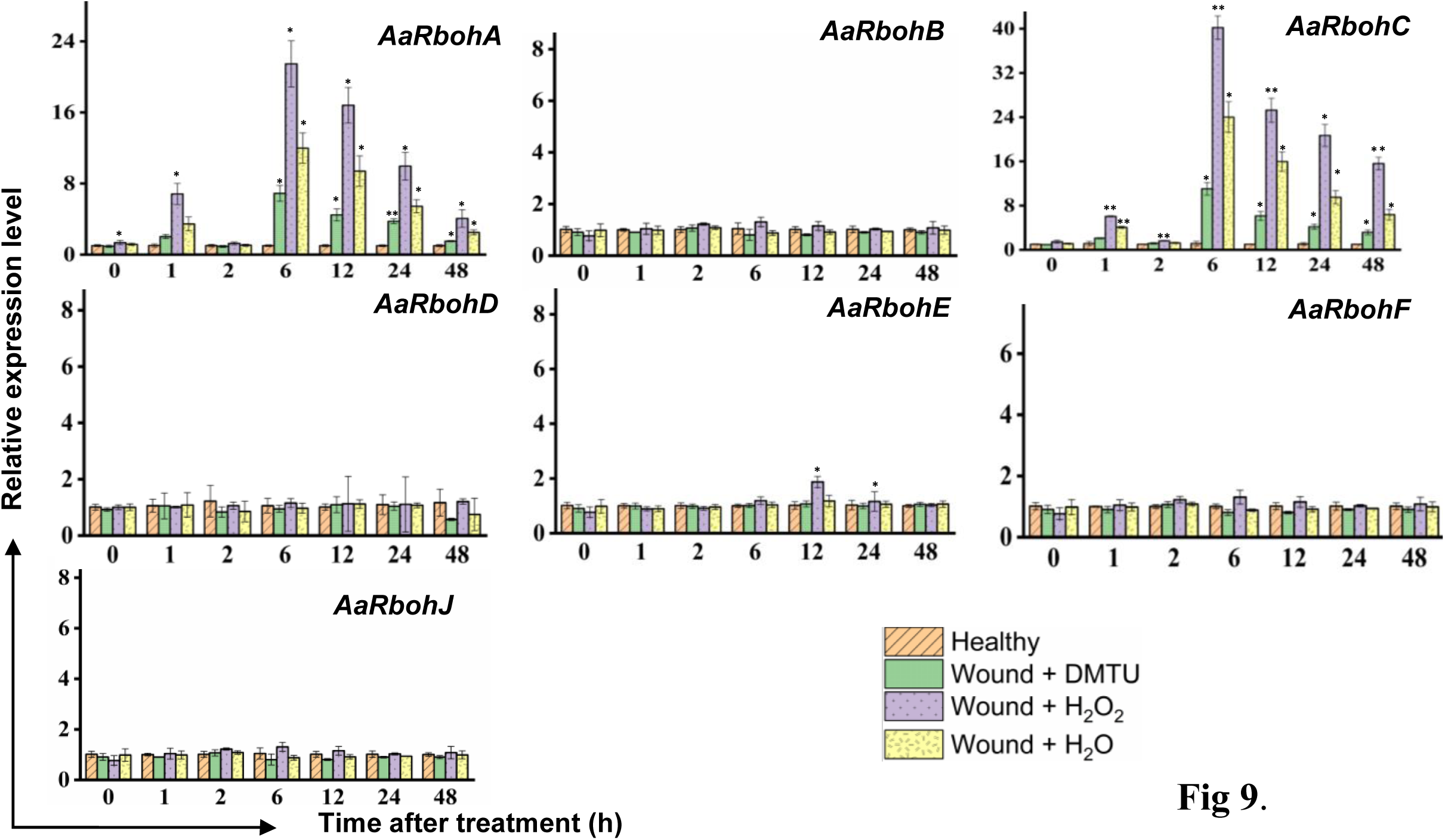
Relative expression levels of *AaRboh* genes in H_2_O_2_-treated stem of *A. agallocha*. The stems were cut, and the apical end of each cut stem was placed in distilled H_2_O, H_2_O_2_, DMTU, respectively as appropriate. The pre-treating solution was thrown away after two h and the stems were left exposed to air. The samples were taken at 0, 1, 2, 6, 12, 24, and 48 h following air exposure. The samples without any treatments are considered as healthy. Asterisks (*) denotes a significant difference compared with healthy samples at 0.05 or **P < 0.01 (Student’s t-test). Data represent means ± SE off three independent experiments.

### 3.9. Transcript abundance of *AaRboh* genes in induced *A. agallocha* tree

The expression of *AaRbohC* and *AaRbohA* elevated in naturally infected wood samples when compared to healthy wood samples (Fig.10). Both genes were found to be significantly upregulated in the range of 22.61 to 76.94-fold in the infected *Aquilaria* wood tissue suggesting their role during stress. This observation also indicates probable role of *AaRbohC* and *AaRbohA* during agarwood induction in *A. agallocha*.

**Fig 10.**
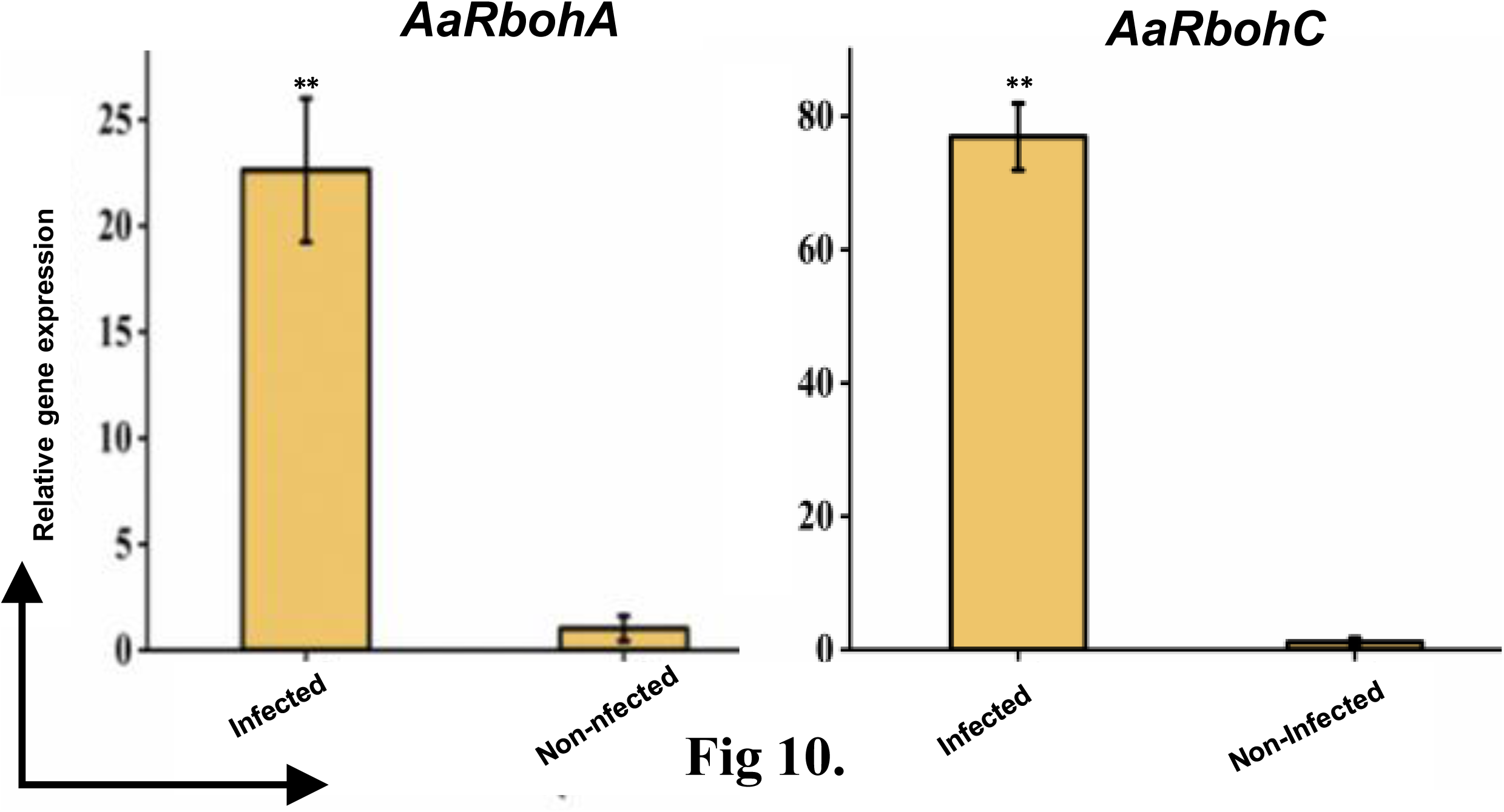
qRT-PCR analysis of 2 *AaRboh* genes. The −2^ΔΔCT^ method was used to determine relative gene expression value. The house keeping gene GAPDH were used to normalized the data. The * symbol indicates transcript levels that differ statistically significantly based on the student t test, and the P value (0.01 < *P < 0.05, **P < 0.01) The mean SE of three technical replicates is used to calculate each expression value. The infected and non-infected plants from Hoollongapar Gibbon Sanctuary.

### 3.10. ROS determination

Treatment with H_2_O_2_ increased endogenous H_2_O_2_ contents both in calli tissues and in actively growing pruned stem tissues. In callus, a transient increase in H_2_O_2_ content was observed after 6 h (3 μmol/g) and then decreased constantly within 48 h (1.3 μmol/g) (Fig. 11A). Moreover, DMTU remarkably decreased the accumulation of H_2_O_2_ and remained constant throughout. As expected, application of DMTU along with H_2_O_2_ decreased the amount of H_2_O_2_ to an average of 0.4 μmol/g in each time period. In wounded stems, H_2_O_2_ concentration reached the first peak at 1 hour (2 μmol/g). Subsequently, started to decrease to the initial level at 2 h and again increased thereafter (Fig. 11B). The maximum H_2_O_2_ production was observed during its second peak at 6 h where intensity was 34 folds of the initial concentration (10 μmol/g). After 24 h of exposure to air, the H_2_O_2_ concentration declined to its lowest level. There was no change in the H_2_O_2_ amount in the healthy stems.

**Fig 11.**
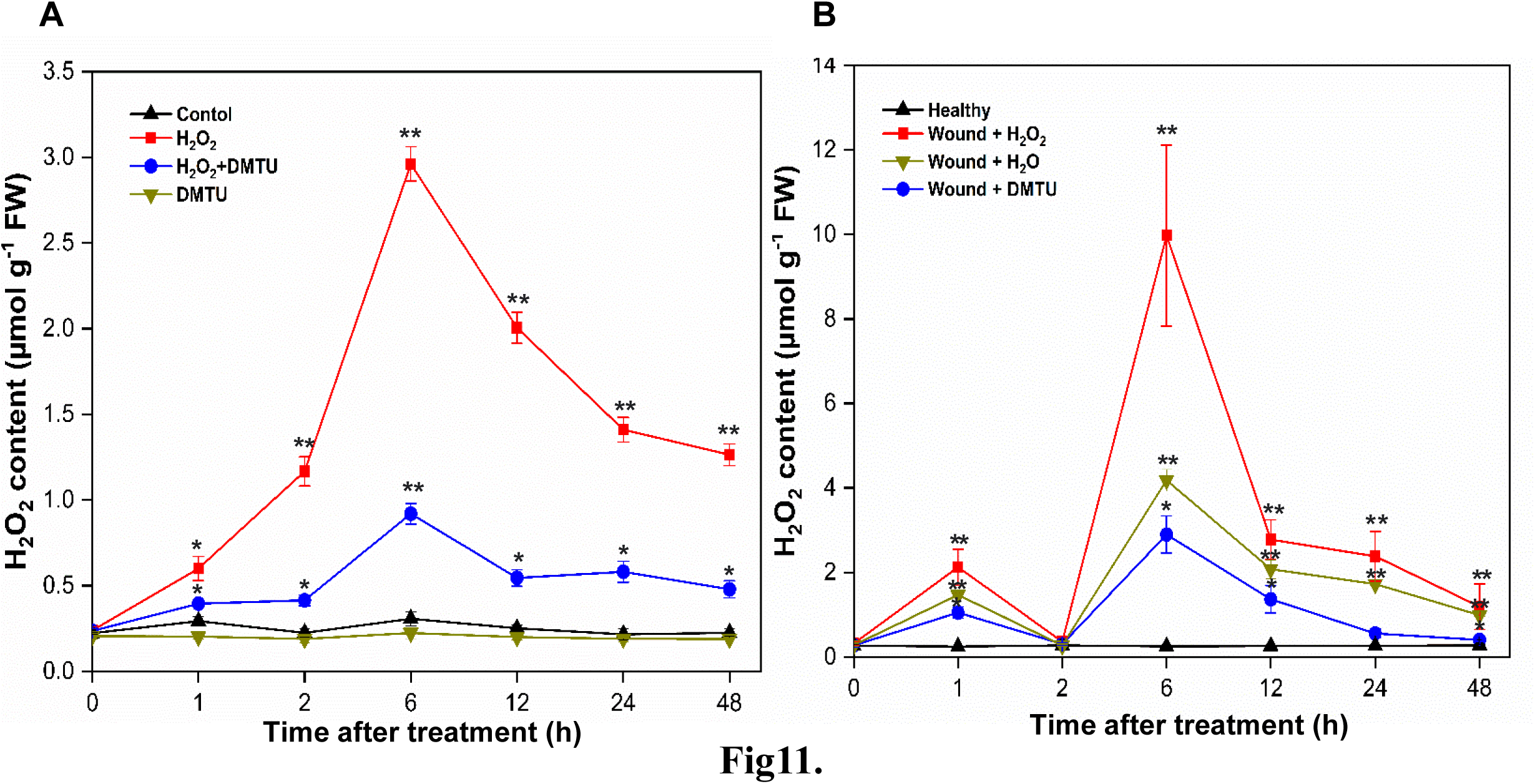
Endogenous H_2_O_2_ content (A) Content of endogenous H_2_O_2_ in calli treated with H_2_O_2_, DMTU and H_2_O_2_ + DMTU, respectively for 0, 1, 2, 6, 12, 24, and 48 h (B) Content of endogenous H_2_O_2_ in the 1-year-old stems after pruning, the cut ends were immersed in distilled H_2_O, H_2_O_2_, DMTU. The pruned stems were exposed to air after 2 h, and the pretreating solution was discarded. The healthy condition indicates the samples without any treatment and served as control. Following air exposure, samples were collected at the 0, 1, 2, 6, 12, 24, and 48 h. Asterisks (*) indicate a statistically significant difference from healthy samples at *P< 0.05 or **P< 0.01 (Student’s t-test). The data represent the means and standard deviations of three independent experiment.

### 3.11 Functional analysis and Protein-protein interaction (PPI)

Gene Ontology based functional prediction revealed the *Aquilaria* Rboh members to be involved in biological processes (BP), molecular function (MF) and cellular component (CC) Supplementary file 7. To understand the physiological and functional interactions of AaRboh proteins, a network model through string database was generated consisting 26 number of nodes and 121 edges (P= 1.0-16) (Fig. 12). Functional information pulled from KEGG database revealed their involvement in MAPK signaling, plant-pathogen interaction and plant hormone signal transduction pathways. In plant pathogen interaction, AaRbohA, AaRbohB, AaRbohC, AaRbohD, and AaRbohE were directly involved and interacted with their functional partners MPK3, CPK28, CDPK1, WRKY33, and EFR. Similarly, in MAPK signaling, all the five Rboh protein were also directly involved and interacted with their partners MKP2, MPK3, WRKY3, ABI1, AB12, and OST1. In hormone transduction, possibly AaRbohA and AaRbohD were involved as they interacted with BR1, AB1, AB2, OST1, ABF2, HAB1, and PP2CA (Fig. 12).

**Fig 12.**
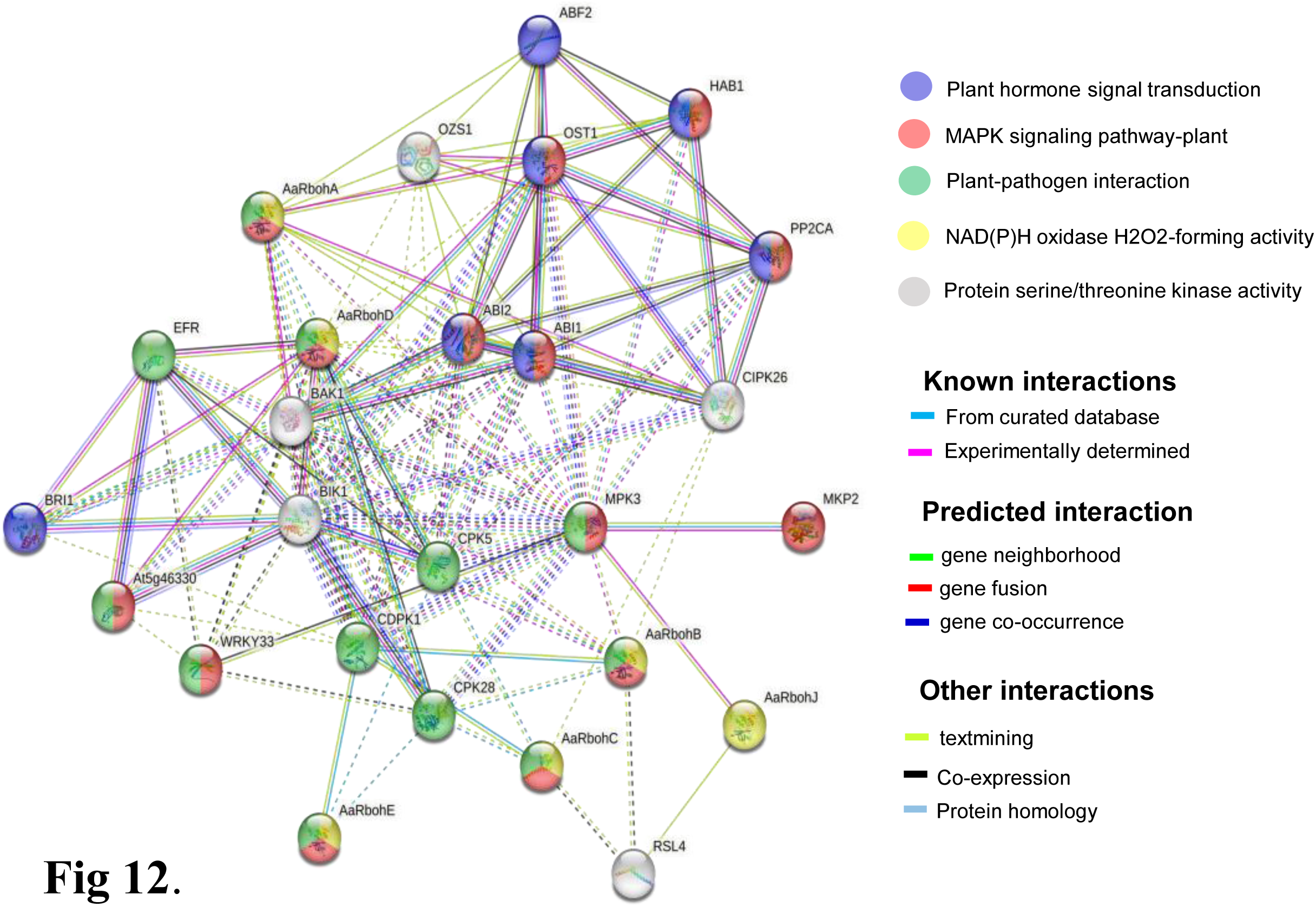
Protein interaction network of AaRboh in *A*. *agallocha* based on *Arabidopsis* orthologues. The potential *AaRboh* with their functional partners in each enriched pathway are displayed in a network model of proteins where the lines of various colours indicate the type of interactions between the potential AaRboh and their functional partners.

## 4. Discussion

### 4.1. Rboh proteins of *A. agallocha* and *A. sinensis*

Rboh protein family have been previously studied in a number of plant species including in *Citrus sinensis* (Zhang et al., 2022), *Capsicum annuum* (Zhang et al., 2021) *Rubus occidentialis*, *Prunus dulsis* (Cheng et al., 2020), *Prunus persica*, Cheng et al., 2019, *Cucumis sativus* (Li et al., 2019), *Jatropha curcas*, *Ricinus communis* Zhao Y and Zou Z 2019, *Fragaria ananassa* cv. Toyonaka (Zhang et al., 2018) and *Vitis vinifera* (Cheng et al., 2013), and interestingly similar number of members have been detected in our study as well i.e., seven in each species of *Aquilaria*. In plants, generation of ROS for development and defence responses is the primary function of Rboh proteins which is also supported by their plasma membrane localization (Sagi and Fluhr 2001). Therefore, recently a number of studies have been carried out to explore their stress biology in many angiosperms (Hu et al., 2018, Kaur and Pati, 2018, Chang et al., 2020; Zhang et al., 2022) The 14 putative Rboh proteins in this study were predicted to be integral components of the cellular membrane, which is in agreement with other plants homologs (Yu et al., 2020), suggesting that they perform similar functions. Location of Rboh proteins other than cell membrane has also been reported indicating their involvement in different function. For instance, FvRbohC in strawberries (Zhang et al., 2018); VvRbohB, VvRbohC2, VvRbohE, and VvRbohF in the grape (Cheng et al., 2013) were located in the thylakoid membrane of chloroplast whereas GaRboh9, GrRboh5, GhRboh2/15, and GbRboh2/15 of cotton (Chang et al., 2020) were restricted to the membrane of mitochondria. Plant *Rboh* have conserved domains in their protein sequences to carry out their primary functions including production of reactive oxygen species during stress and regulation of Ca^2+^ channels and downstream activation (Yu et al., 2020; Torres et al., 2005). *Aquilaria Rboh* possessed evolutionary conserved domains, with an additional TM domain possibly for ferric reductase activity and two Ca^2+^-binding structural motifs EF-hand, which directly indicate that the ROS produced by the Rboh protein is part of a cellular signalling network in plants. This crucial motif (EF hands) is absent from their mammalian homolog pg91phox, demonstrating a distinct mechanism for the generation of ROS in plants and animals (Torres et al., 2005). Presence of these domains in the putative Rboh proteins of both *Aquilaria* spp might be due to their association with stress induced ROS generation.

### 4.2. Phylogenetic relation of Aquilaria Rboh proteins with other plants

*Aquilaria* Rboh proteins clustered into five groups, with closer evolutionary relationship to soybean and potato, followed by *Arabidopsis* in the phylogenetic tree. The result is consistent with previous reports illustrated in other plants (Zhang et al., 2018, Zhang et al., 2021, and Zhang et al., 2022). Therefore, homologs clustered in the same group may possess close functional relation (Ito et al., 2017).Gene duplication is caused by three mechanisms, i.e., whole-genome duplication (WGD), segmental duplication, and tandem duplication. (Zhang et al., 2021). Previous studies have identified that WGD and segmental duplication are the major driving force for gene duplications of the *Rboh* gene family in plant species such as *Brassica rapa* (Li et al., 2019)*, Musa acuminate* (Ying et al., 2020), and *Gossypium hirsutum* (Wang et al., 2020). In *A. thaliana*, expansion of *Rboh* genes were due to WGD and local duplication (Zhou et al., 2019), while in *G. max* and *C. annuum*, segmental duplication was the major events for expansion (Liu et al., 2019, Zhang et al., 2021). Interestingly, during the domestication of *Malus domestica*, *Rboh* genes were duplicated by both WGD and segmental duplication events (Cheng et al., 2020). In this study, we found one segmental duplication event in *A. sinensis*, whereas no duplication events were observed in the *A. agallocha* genome. The paralogous gene pair had a Ka/Ks value of 0.54, which was substantially lower than 1, indicating that these genes underwent strong purifying selection. In addition, the decreased number of Rboh proteins in the both the *Aquilaria* genome compared to *Arabidopsis* implies that probable loss of function event had occurred during the evolution of the *Aquilaria* genome, or functional redundancy may also be the cause of gene loss events (Adriana et al., 2015, Martin and Schnarrenberger et al., 1997).

### 4.3. Functional role of *AaRboh* genes upon stress in callus

The cis-regulatory elements of 14 putative *Rboh*s supported an earlier study which demonstrated their role in stress, hormonal responses, and development processes (Zhang et al 2022) and the results are consistent with *O. sativa* and *A. thaliana* (Kaur et al., 2016). In addition, evidences elucidated from KEGG pathway analysis and interaction network also support their involvement in plant-pathogen interaction, hormone, and MAPK signalling. In other plants, literature demonstrated evidences for their involvement in development (Potocký et al., 2007; Marino et al., 2012), stress (Si et al., 2010; Yu et al., 2020; Huang et al.,2021; Torres et al., 2002; Liu et al., 2019), and hormone signalling (Jakubowicz et al., 2010). Moreover, Rboh proteins of *H*. *vulgare*, *G. barbadense*, *Z. jujube*, *J. curcas* and *M. sativa* have been shown to participate in ROS accumulation and provide resistance against infecting plant pathogens (Trujillo et al., 2006, Zhao et al., 2019, Cheng et al 2020). Therefore, for functional validation of *Aquilaria* Rboh members, we extended our study to analyse the expression level of these members in stress induced callus tissues and wounded stem through qRT-PCR, and to estimate ROS after treatment. Elevation in the expression of the two members viz., *AaRbohA* and *AaRbohC*, and accumulation of maximum ROS inside both callus and stem after 6 h of treatment indicates their association with ROS production upon stress. Additionally, elevation of *AaRbohB* and *AaRbohE* only in callus further indicates their contribution in generation of ROS specifically in callus tissue. The genes *AsRbohA-C* also could accumulate endogenous H_2_O_2_ in *A. sinensis* callus upon salt stress (Wang et al., ^2^01^8^). In other plant such as *C*. *annum* L. cold, drought and salt stresses mediate high amount of endogenous H_2_O_2_ (Zhang et al., 2021). Besides H_2_O_2_ and salt methyl jasmonate also upregulated expression of *Rboh* genes considerably in calli of *A. sinensis* (Xu et al., 2013).

The same have also been observed in *Nitraria tangutorum* (Yang et al., 2012). Therefore, it can be stated that upon stress in callus of *Aquilaria, Rboh* genes generate ROS which can further lead to the accumulation of sesquiterpenoids and 2-(2-phenylethyl)chromones as a defense response (Zhang et al., 2014, Xu et al., 2016, Lv et al. in 2019). The ROS generated may likely act as signaling agent for modulation of genes involved in biosynthesis of secondary metabolites including sesquiterpenoids and 2-(2-phenylethyl)chromones in *Aquilaria* callus tissues.

### 4.4. Involvement of *AaRbohA* and *AaRbohC* during agarwood formation in *A. agallocha*

It is a widely accepted fact that agarwood in *Aquilaria* plant species is produced naturally due to plant-fungal interaction (Ahmead and Kulkarni 2017, Gao et al., 2019, Haung et al., 2022). Wound induced signalling mechanism and the mitogen-activated protein kinase (MAPK) signaling pathway has been proposed to be associated with secondary metabolite biosynthesis leading to agarwood formation in *A*. *sinensis* (Tan et al., 2019). Recent research on similar line suggested ROS mediated activation of MAPK pathway during plant-pathogen interaction (Pacheco-Trejo et al., 2022; Chuang et al., 2022). Interestingly, in other plants as well, *Rboh* have been shown to be involved in responses to microbial infection (Morales et al., 2016, Chaung et al., 2022). Therefore, we further extended our study to estimate the abundance of *AaRboh* genes in agarwood producing naturally infected *A. agallocha* tree, and observed that the same two members i.e., *AaRbohA* and *AaRbohC* significantly upregulated when compared to healthy (non-infected plant). This indicates their involvement in production of ROS in the tissues post fungal infection in *A. agallocha* tree leading to H_2_O_2_ burst (Rejeb et al., 2015, Wang et al., 2018). This probably induces the loop of plant hormone biosynthesis including MeJA (Xu et al., 2013) leading to modulation of genes involved in biosynthesis of metabolites required for defence response (Lv et al., 2019, Das et al., 2023) through transcription factors such as MYC2 and WRKY (Xu et al., 2016, 2017; 2020), and in due course agarwood formation occurs. Interaction of *AaRbohA* and *AaRbohC* with WRKY in our constructed network also supports the observation. In addition, presence of high number of defense and stress related cis-acting elements in the promoters, and their association with MAPK signaling, plant-pathogen interaction and plant hormone signal also support our prediction that these two members through ROS generation play a crucial role during microbial invasion and during agarwood formation *in A. agallocha*.

## 5. Conclusion

In this study, we comprehensively identified seven putative *Rboh* genes each in the two species of *Aquilaria* i.e., *A. agallocha* and *A. sinensis* which were clustered in five groups with their close homologs of other plants. The exon-intron structure and distribution of motif pattern showed the divergence and conservation of the members. Presence of orthologous gene pairs with *Rboh* genes of other plant indicated their evolutionary and close functional relation. Evidences generated from the qRT-PCR analysis coupled with ROS estimation suggest that the proteins encoding *AaRbohA* and *AaRbohC* are involved in ROS generation; most probably during fungal invasion they may serve as signaling molecule during various stages of agarwood formation in *A. agallocha*. The findings of this study contribute to a fundamental understanding of the molecular basis of the *Rboh* gene family in the *Aquilaria* genome and establish a foundation for future research on this gene family. This knowledge will further enhance our understanding of the ROS network and its influence on agarwood formation.

## Supporting information

Supplentary file 1: Gene-specific primers for qRT-PCR analysis used in this study

Supplentary file 2: Domain organization of 14 Rboh genes from A. agallocha and.A. sinensis

Supplentary file 3: Cis regulatory elements in the 2000 promoter region of 7 AaRboh and 7 AsRboh

Supplentary file 4: Orthologous gene pairs

Supplentary file 5: Duplicated gene pair

Supplentary file 6: Log fold value of transcripts of AaRboh

Supplentary file 6: Gene ontology and KEGG functional prediction of AaRboh proteins

## Data availability

The data supporting the finding of this study is provided in the manuscript and itssupplementary material.

## Ethics approval

Not applicable

### Acknowledgement

The authors are indebted to Gauhati University for providing the technical facility. The authors also acknowledge Department of Biotechnology (DBT), Government of India for providing the financial aid.

### Abbreviations

*Rboh*: Respiratory burst oxidase homolog
AaRboh: Aquilaria agallocha Respiratory burst oxidase homolog
ROS: Reactive Oxygen Species DMTU Dimethylthiourea
HMM: Hidden Markov Model
CDS: Coding Sequences
GO: Gene ontology
KEGG: Kyoto Encyclopaedia of Genes and Genomes database

## Supplementary files

**Supplentary file 1**: Gene-specific primers for qRT-PCR analysis used in this study

**Supplementary file 2.** Domain organization of 14 *Rboh* genes from *<u>A. agallocha</u>* and*.A. sinensis* The characteristics domains are displayed based upon results of putative *Rboh* SMART tool search (http://smart.embl-heidelberg.de/)). Blue rectangular boxes represent trans-membrane regions and small pink boxes represents low complexity regions.

**Supplimentary file** 3: Cis regulatory elements in the 2000 promoter region of 7 AaRboh and 7 *AsRboh*

**Supplinatary file 4**: Orthologous gene pairs

**Supplimentary file 5(A**): Duplicated gene pair

**Supplimentary file 5(B**): Ka/Ks value calculation

**Supplimentary file 6(A):** Log fold value of transcripts of *AaRboh*

**Supplimentary file 6(B**): Log fold value of transcripts of AsRboh

**Supplimentary file 7:** Gene ontology and KEGG functional prediction of AaRboh proteins

## Notes

### Competing Interest Statement

The authors have declared no competing interest.

