## Supplementary figures and images for "Genome-wide analysis of Respiratory burst oxidase homolog (*Rboh*) genes in *Aquilaria* species and its association with agarwood formation"

### Supplentary file 2: Domain organization of 14 Rboh genes from A. agallocha and.A. sinensis

#### Slide 1
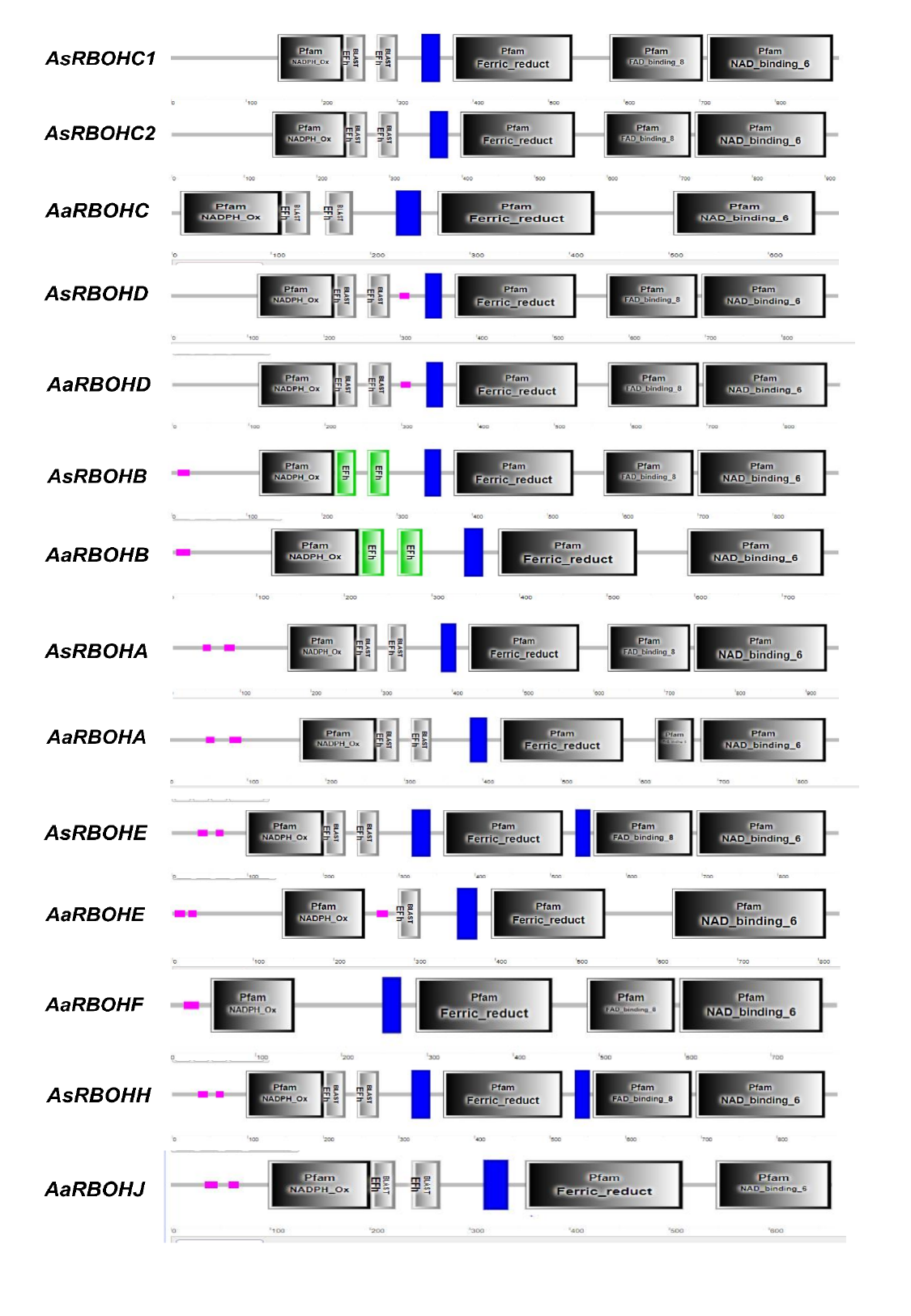

#
